# Sulfur Isotope Effects of Dissimilatory Sulfite Reductase

**DOI:** 10.1101/023002

**Authors:** William D. Leavitt, Alexander S. Bradley, André A. Santos, Inês A.C. Pereira, David T. Johnston

**Affiliations:** Department of Earth & Planetary Sciences, Harvard University, Cambridge, MA, USA; Department of Earth and Planetary Sciences, Washington University in St. Louis, MO, USA; Instituto de Tecnologia Química e Biológica António Xavier, Universidade Nova de Lisboa, Oeiras, Portugal

## Abstract

The precise interpretation of environmental sulfur isotope records requires a quantitative understanding of the biochemical controls on sulfur isotope fractionation by the principle isotope-fractionating process within the S cycle, microbial sulfate reduction (MSR). Here we provide the only direct observation of the major (^34^S/^32^S) and minor (^33^S/^32^S, ^36^S/^32^S) sulfur isotope fractionations imparted by a central enzyme in the energy metabolism of sulfate reducers, dissimilatory sulfite reductase (DsrAB). Results from in vitro sulfite reduction experiments allow us to calculate the in vitro DsrAB isotope effect in ^34^S/^32^S (hereafter, ^34^ε_DsrAB_) to be 15.3±2‰, 2σ.The accompanying minor isotope effect in ^33^S, described as ^33^λ_DsrAB_, is calculated to be 0.5150±0.0012, 2σ. These observations facilitate a rigorous evaluation of the isotopic fractionation associated with the dissimilatory MSR pathway, as well as of the environmental variables that govern the overall magnitude of fractionation by natural communities of sulfate reducers. The isotope effect induced by DsrAB upon sulfite reduction is a factor of 0.3 to 0.6 times prior indirect estimates, which have ranged from 25 to 53‰ in ^34^ε_DsrAB_. The minor isotope fractionation observed from DsrAB is consistent with a kinetic or equilibrium effect. Our in vitro constraints on the magnitude of ^34^ε_DsrAB_ is similar to the median value of experimental observations compiled from all known published work, where ^34^ε_r-p_ = 16.1‰ (*r – p* indicates reactant versus product, n = 648). This value closely matches those of MSR operating at high sulfate reduction rates in both laboratory chemostat experiments (^34^ε_SO4-H2S_ = 17.3±1.5‰) and in modern marine sediments (^34^ε_SO4-H2S_ = 17.3±3.8‰). Targeting the direct isotopic consequences of a specific enzymatic processes is a fundamental step toward a biochemical foundation for reinterpreting the biogeochemical and geobiological sulfur isotope records in modern and ancient environments.

## Introduction

Microbial sulfate reduction provides a critical link between Earth’s exogenic sulfur, carbon, iron and oxygen cycles (Thode et al., 1961; Canfield, 2001a; Garrels and Lerman, 1981; Holland, 1973). This metabolism is comprised of a set of enzymes working in concert to reduce sulfate 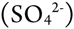 to sulfide (H_2_S) (Pereira et al., 2011; Peck, 1961) (Figure 1). During this transformation, MSR generates ^34^S/^32^S, ^33^S/^32^S, ^36^S/^32^S, ^18^O/^16^O and ^17^O/^16^O stable isotope fractionations (Harrison and Thode, 1958; Kaplan and Rittenberg, 1964; Chambers et al., 1975; Kemp and Thode, 1968; Canfield, 2001b; Goldhaber and Kaplan, 1975; Leavitt et al., 2013; Sim et al., 2011b; Fritz et al., 1989), the biochemical source of which is unclear (Chambers and Trudinger, 1979). To construct a biochemically constrained perspective of sulfur isotope fractionations during MSR requires we quantify how material moves through the metabolic network, and gain an understanding of the isotope effect(s) associated with each constituent enzymatic step (Hayes, 2001).

**Figure 1:**
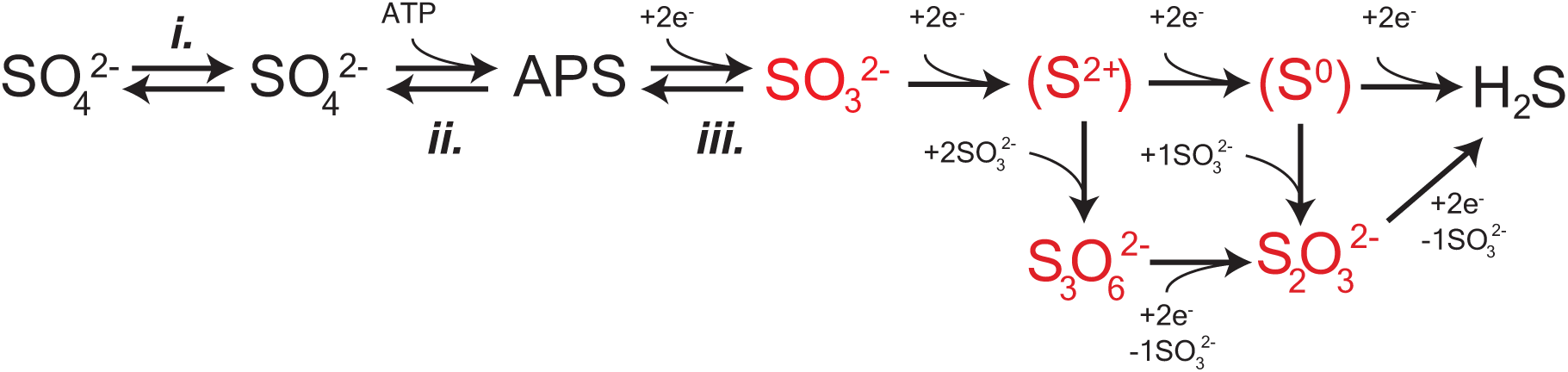
A schematic capturing the central role of DsrAB in MSR. The in vivo dissimilatory sulfate reduction pathway, where red highlighted steps represent sulfite reduction by DsrAB in the absence of DsrC, as targeted here in vitro. The constituent steps of MSR relevant to S isotope fractionation are likely APS reduction to sulfite by APSr, sulfite reduction (the subject of this study) by DsrAB, and the terminal production of sulfide by DsrC/DsrMKJOP. The pathway is described in detail in the text.

The enzyme catalyzed reaction network of MSR is represented in Figure 1. Sulfate is first imported into the cytoplasm by a variety of transporters (Cypionka, 1994; Piłsyk and Paszewski, 2009) (Figure 1), and subsequently activated to a high-energy intermediate, adenosine 5’-phosphosulfate (APS). The latter reaction generates pyrophosphate (PPi) at the expense of ATP by the enzyme sulfate-adenylyl transferase (Sat) (Peck, 1962). APS is reduced to sulfite 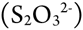 through a two-electron transfer by the soluble cytoplasmic enzyme APS oxidoreductase (ApsR) (Peck, 1959), which is linked to energy conservation by the membrane-bound complex QmoABC (Pires et al., 2003). APS reduction is highly reversible, depending on the in vivo or in vitro conditions (Peck, 1960). Sulfite has several potential fates. Sulfite can either be re-oxidized to sulfate (directly or via APS) or further reduced to sulfide by DsrAB with the involvement of DsrC (Oliveira et al., 2008b). A critical step is during the reduction of sulfite when it binds the iron of the siroheme in the DsrAB active site. The subsequent reduction occurs via electron transfer from an adjacent Fe-S cluster (Oliveira et al., 2008a; 2008b; Parey et al., 2010). In vivo, DsrAB has been proposed to generate intermediate valence sulfur, which is then bound to DsrC and converted to sulfide via DsrC/MK (Oliveira et al., 2008b; Venceslau et al., 2013; 2014). Sulfide then leaves the cell by diffusion (as H_2_S) or through anion transport (as HS^-^ or S^2-^). In instances when DsrC is unavailable (e.g. when DsrAB is pure in vitro) or limiting (e.g. intracellular sulfite is in excess of reduced DsrC), intermediates such as thiosulfate 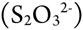 may become important, likely due to the reaction of sulfite with sulfide (in vivo) or the partially reduced sulfur from DsrAB (in vitro) (Drake and Akagi, 1976; Chambers and Trudinger, 1975; Drake and Akagi, 1978; Kim and Akagi, 1985; Drake and Akagi, 1977). A few examples exist where thiosulfate is a key component in closing S mass balance during in vivo MSR (Sass et al., 1992; Price et al., 2014; Leavitt et al., 2014), and in one instance trithionate is observed 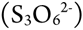 (Sass et al., 1992), though it is not clear in this case it is a physiological product. Under these conditions, accumulation and excretion of such compounds as thiosulfate may be important (Bradley et al., 2011). It is within this broader biochemical and physiological context that we examine the isotopic consequences of sulfite reduction by DsrAB, which as outlined above, is central to the biochemistry of dissimilatory sulfate reduction.

Reduction of sulfite by DsrAB breaks three of the four S-O bonds in the original sulfate (Venceslau et al., 2014). As such, the isotope effect of DsrAB likely plays a significant role in setting the overall fractionations observed from MSR (Harrison and Thode, 1958; Rees, 1973; Brunner and Bernasconi, 2005; Farquhar et al., 2003). Measured enzyme-specific isotope effects are lacking for MSR and the S cycle in general. Such information has been transformative for the study of other biogeochemical elements like carbon. For example, experimental work quantifying the ^13^C/^12^C effect of RuBisCO (Park and Epstein, 1960; Farquhar et al., 1982; Tcherkez et al., 2006), the core enzyme in carbon fixation, has greatly advanced the applicability of carbon isotope biogeochemistry. More specifically, understanding the fractionation associated with RuBisCO allowed greater insight into modern (Hayes, 1993) and ancient (Hayes et al., 1999) carbon cycling, and facilitated a better understanding of primary productivity in the both modern (Laws et al., 1995) and ancient (Pagani et al., 2009) oceans. Similar approaches have also proven greatly informative in studies of methane production (Scheller et al., 2013), nitrate assimilation (Karsh et al., 2012), and nitrogen fixation (Sra et al., 2004). With these studies as a guide, we look to further unlock the sulfur cycle through targeting a key microbial sulfate reduction enzyme, DsrAB.

To close the knowledge gap between whole-cell observations and enzyme-catalyzed reactions, as well as to turn natural isotope records into catalogues of environmental information, we conduct the first enzyme-specific sulfur isotope experiments. Here we report the sulfur isotope fractionation factors associated with in vitro sulfite reduction by the dissimilatory sulfite reductase enzyme (DsrAB). Using these results and a new mathematical model, we are able to place improved constraints on the root of sulfur isotope fractionation during MSR. This refines our understanding of the predominant biological process responsible for generating environmental S isotope records throughout geological history.

## Experimental Methods Summary

We conducted a series of closed system in vitro sulfite reduction experiments with purified DsrAB from *Desulfovibrio vulgaris* str. Hildenborough (DSM 644) and *Archaeoglobus fulgidus*. These enzymes are structurally similar and evolutionarily related (Parey et al., 2013), and we chose them to attempt to determine conservation of isotope fractionation in *D. vulgaris* and *A. fulgidus* DsrAB. The complete isolation and purification details are available in the Appendix.

DsrAB experiments were conducted in vitro under strictly anoxic conditions with H_2_, [NiFe] hydrogenase, and methyl viologen as the electron donation system. Key considerations in experimental design are: (i) to provide enough sulfur at each time point for isotopic characterization of residual reactant and products; (ii) to provide the proper reaction conditions to allow for optimal DsrAB activity (pH = 7.1, T = 20° or 31°C); (iii) to ensure hydrogenase activity is not inhibited by the experimental pH (optimum at pH 7.5, activity significantly depleted below pH 6.5, so we chose pH 7.1, to account for optima of both DsrAB and [NiFe]-hydrogenase); and finally (iv) to ensure the sulfite to hydrogen ratio favors sulfite reduction. Experiments setup is detailed in the Appendix.

Each experiment was performed in duplicate and sampled as sulfite was consumed (reaction progress tracked as *f*_SO3_, equivalent to the fraction of remaining sulfite). The reaction consumed sulfite to form products thiosulfate and trithionate, with no detectable sulfide. Thiosulfate and trithionate concentrations were quantified following published cyanolysis protocols (Kelly and Wood, 1994), where we used a modified ‘Fuschin’ method (Grant, 1947) to quantify sulfite and a modified Cline method (Cline, 1969) to measure sulfide. All quantification and experimental methods are fully detailed in the Appendix. In addition to concentrations, we measured the major and minor sulfur isotopic compositions of three operationally defined and precipitated pools: sulfite (both initial and residual reactant), product sulfonate (from trithionate or thiosulfate) and the ‘reduced sulfur’ reservoirs (central and terminal sulfurs in trithionate and thiosulfate, respectively). Complete IUPAC definitions of each S reservoir, along with all isotopic measurement methods and error propagation calculations are fully articulated in the Appendix.

## Isotope Notation

The variability in ^34^S of a measured pool is reported in standard delta notation (for instance δ^34^S, in ‰ units), where ^34^S/^32^S of the sample is the relative difference from a standard (Hayes, 1983), and is reported as the isotopic offset between two measured pools of sulfur, ^34^ε (=10^3^x(^34^α-1)), still in ‰ units. Fractionation factors (α’s and associated ε’s) are annotated with a subscript to denote the process of interest or pools being related, such as ^34^ε_DsrAB_, ^34^ε_MSR_, ^34^ε_*r-p*_ or ^34^ε_*SO4-H2S*_. The same nomenclature convention is followed when a minor isotope, ^33^S, is included. The only exception is the addition of one new term, ^33^λ, which is approximately the slope of a line on a plot of δ^33^S versus δ^34^S (Farquhar et al., 2003; Miller, 2002), but can be simply interpreted as a measure of mass-dependent minor isotope fractionation. Mathematical definitions are provided below.

## Fractionation modeling

Calculation of the isotopic fractionation imposed by the reduction of sulfite through DsrAB requires tracking the concentration of the reactant, accumulation of the products, and determining the isotopic composition of all as the reaction progressed. This necessitates the application of a closed-system model in order to calculate fractionation factors. Determining the intrinsic isotope effect associated with a closed system reaction can be approached in a number of ways. Normally, in a system where one reactant is consumed in order to generate a single product, a Rayleigh model is employed (Mariotti et al., 1981; Nakai and Jensen, 1964). This approach assumes that the reaction of interest is unidirectional, generates only one product, and that the fractionation factor is invariant throughout the reaction. In this case, the isotope effect is calculated as a function of the isotope ratio, *R*, of the starting composition (*R*_*ao*_) and evolving product pool (*R*_*p*_, defined below), equal to the mass balance on sulfite:

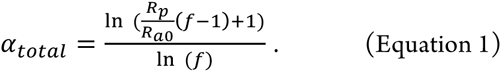

In this solution, *f* tracks the fractional amount of reactant remaining 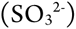. For our experiments, we define *f*_SO3_:

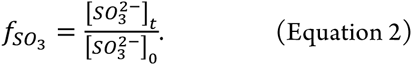

In the specific case of our experiments and the reduction of sulfite by DsrAB, however, the standard closed-system isotope distillation models (equation 1) requires expansion. Recall that the in vitro reaction involves the accumulation of two products (trithionate and thiosulfate). Each of these products further contains sulfur moieties in more than one oxidation state (Kobayashi et al., 1974; Drake and Akagi, 1977; 1978; Suh and Akagi, 1969; Drake and Akagi, 1976). This means that, rather than R_p_ being the isotope composition of a single product pool, we define it as the mass-weighted sum of the oxidized (*R*_ox_) and reduced (*R*_red_) products in trithionate and thiosulfate:

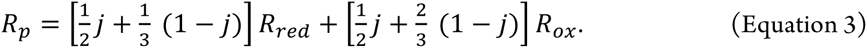

Here the reduced and oxidized pools are the operationally defined reservoirs discussed above. In the mass balance accounting equation we introduce a term to quantify the ratio of products, *j* (Figure 2). The concentrations of sulfite, trithionate, and thiosulfate were measured at each time-point, ensuring the closure of mass balance and validating the use of a relative mass term. The *j* term is thus the fraction of products residing in thiosulfate:

**Figure 2:**
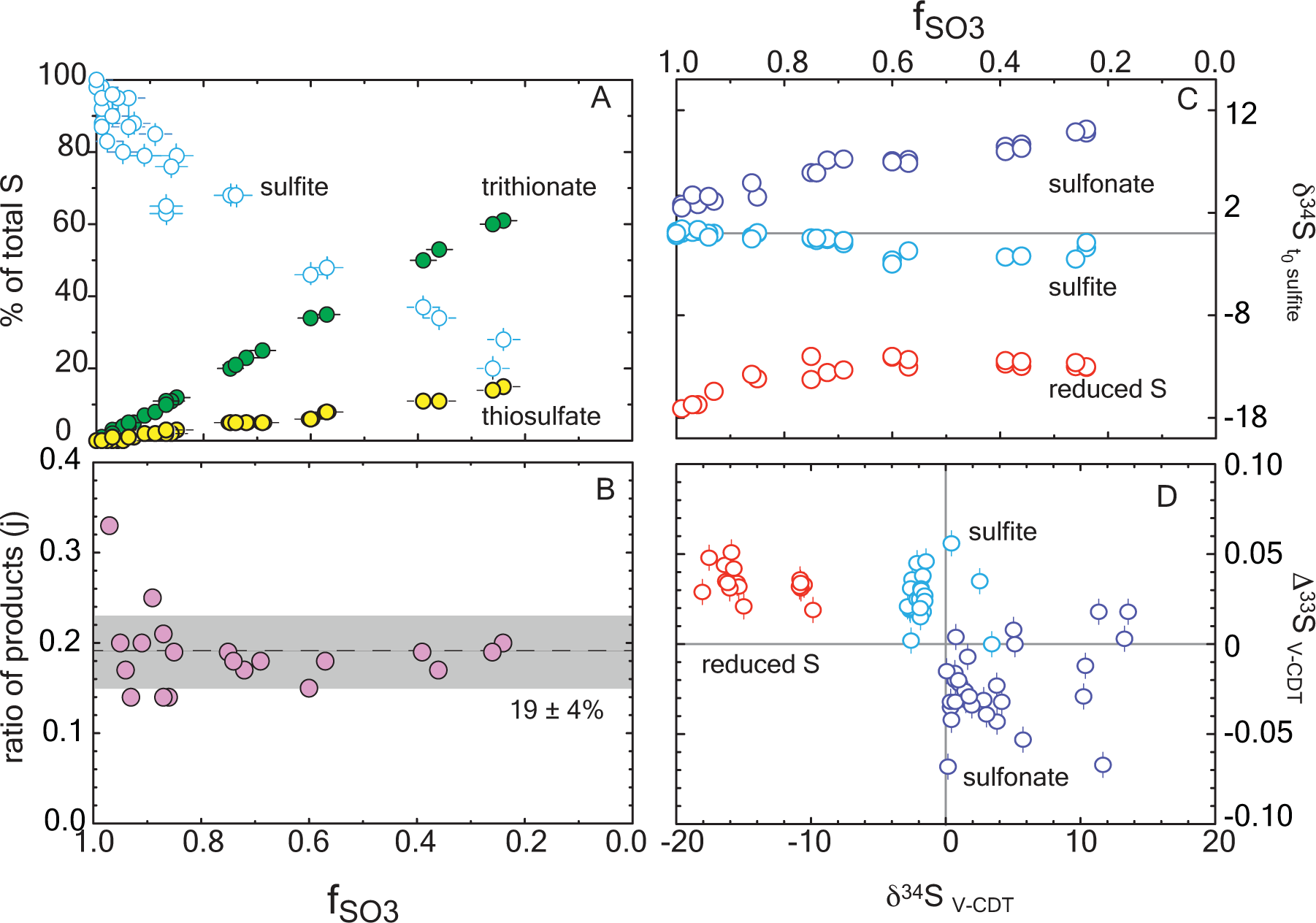
Reaction progress during sulfite reduction with *D. vulgaris* DsrAB in vitro. Errors are included in all measurements (2σ) and are smaller than the symbol if not seen. (A) The mol fraction of sulfur in each sulfur product pool as a function of reaction progress, *f*_SO3_. Mass balance conservation is discussed in the text. (B) The ratio of products at each time point, demonstrating the constancy of the reaction scheme (denoted as *j* in the model). This is the ratio of the slopes of the products from A. (C) Major isotope data for each operationally defined sulfur pool as a function of reaction progress, and normalized to the initial sulfite composition. (D) A triple isotope cross plot of the data presented in frame C, normalized to V-CDT.

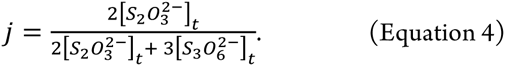

For isotopic measurements we quantitatively separated the oxidized moieties from trithionate and thiosulfate from the partially reduced moieties of both products. There were no available methods to separate trithionate and thiosulfate and isolate each S site within those products (a target for future work). We then measured the isotopic compositions of the pooled oxidized and pooled reduced products. As the goal is to identify the fractionation between the residual sulfite and either the oxidized (^3x^α_ox_) or reduced (^3x^α_red_) moieties in trithionate and thiosulfate, we present the general equation, (^3x^α_z_):

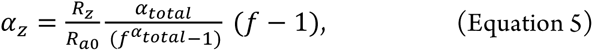

where *z* is either *ox* or *red*. This solution is then translated into standard ^3x^ε notation. Fractionation factors are then related in triple isotope space with:

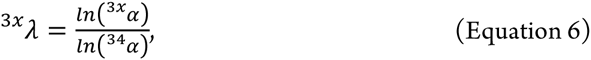

a term which finds common application in mass-dependent studies (Young et al., 2002; Farquhar et al., 2003). Finally, we note the models assumptions: 1) sulfite and its isomers carry the same isotopic composition as each other, 2) the isotopic composition of the sulfonate groups (in trithionate and thiosulfate) are isotopically identical, and 3) similar to 2, the isotopic composition of reduced sulfur in trithionate and thiosulfate are isotopically identical.

The complexity added above in equation 6 allows for numerous products for a given reaction, but still assumes that the fractionation factors involved are static over the time series of the experiment (*f*_SO3_) and that there is only one reaction present. If this is true, then the model prediction will match the observation over all values of *f*_SO3_. Although we observed a statistically invariant ratio of thiosulfate to trithionate production throughout the reaction (*j* in Figure 2), suggesting a static set of reactions through the entire experiment, it appears that the net fractionation factor was indeed time-dependent. In the event of an evolving α, the fractionation factor early in the experiment, where the concentration of products remains low, most closely approximates the isotope effect of DsrAB solely reducing sulfite. We explore this time-dependence further in the Appendix for all sampled points on the reaction progress coordinate (*f*_SO3_). Thus, for extracting the intrinsic isotope effect associated with enzymatic reduction of sulfite, we focused on data where *f*_SO3_ > 0.85. To do so, we have used our modified Rayleigh-type isotope distillation model in which we account for the production of reduced and oxidized sulfur within aqueous products trithionate and thiosulfate. Procedures for error propagation associated with these calculations are described in the Appendix.

## Results

We tracked all S pools at each time point. Mass balance was satisfied within ±10% of the initially provided sulfite in every experiment, and within ±5% in 27 of 33 experiments (Figure 2a). The majority of this variance is due to analytical error in the sulfite quantifications. In experiments with the *D. vulgaris* DsrAB, the products were generated with a mean of 19% of the product sulfur forming thiosulfate and the remainder accumulating as trithionate (Figure 2b). This is consistent with previous reports (Drake and Akagi, 1976; 1978), and expected given the absence of active DsrC in these experiments. Some inactive DsrC does accompany the *D. vulgaris* DsrAB during isolation and purification (Oliveira et al., 2008a; 2008b; 2011), however there is no means to recycle this component, and as such, it is not a functional part of the experiment. Therefore, the in vitro sulfite reduction reactions produce thionates rather than sulfide. In our experiments, sulfite was always in excess and never became limiting.

To extend our studies to a different taxonomic form of the enzyme, we also experimented on the DsrAB from the thermophilic archaeon *A. fulgidus*. This enzyme operates at higher temperature and lacks DsrC in the complex (Schiffer et al., 2008). *A. fulgidus* DsrAB experiments were conducted with 15 mM initial sulfite and at 65°C, where they showed consistent loss of sulfite and accumulation of products between replicates at each time point. Unlike *D. vulgaris* DsrAB experiments, however, only small quantities of product were generated. From these experiments we were able to resolve a complete sample set (i.e. sulfite, sulfonate and reduced S) from one time-point and partial sets from another (i.e. sulfite and sulfonate). Special efforts were made to correct data available on *A. fulgidus* experiments (see the Appendix). The results using DsrAB from *A. fulgidus* are consistent with those of *D. vulgaris*, but with a large calculated uncertainty. We therefore focus our interpretations on the results from the *D. vulgaris* experiments.

We use the concentrations of sulfite, trithionate, and thiosulfate, as well as the isotopic compositions of each operationally defined product to solve for the fractionations associated with DsrAB. The calculated ^34^ε_DsrAB_ for sulfite reduction by the *D. vulgaris* DsrAB is 15.3±2.0 ‰ (2σ, Figure 3), where the concurrent fractionation associated with the generation of the sulfonate is -3.2±0.8‰ (2σ). The ^34^ε_DsrAB_ from *A. fulgidus* is generally consistent with the *D. vulgaris* experiments, yielding a reductive fractionation of 16‰ (2σ from 22 to 12‰) at 65°C. Large and asymmetric errors on the *A. fulgidus* data are the result of exceptionally small sample sizes, which also precluded the collection of ^33^S data (see the Appendix). Together, these experiments demonstrate a broad consistency in fractionation by DsrAB over a wide range of temperatures (20 and 30°C for *D. vulgaris*, and 65°C for *A. fulgidis*) and across two Domains of life.

**Figure 3:**
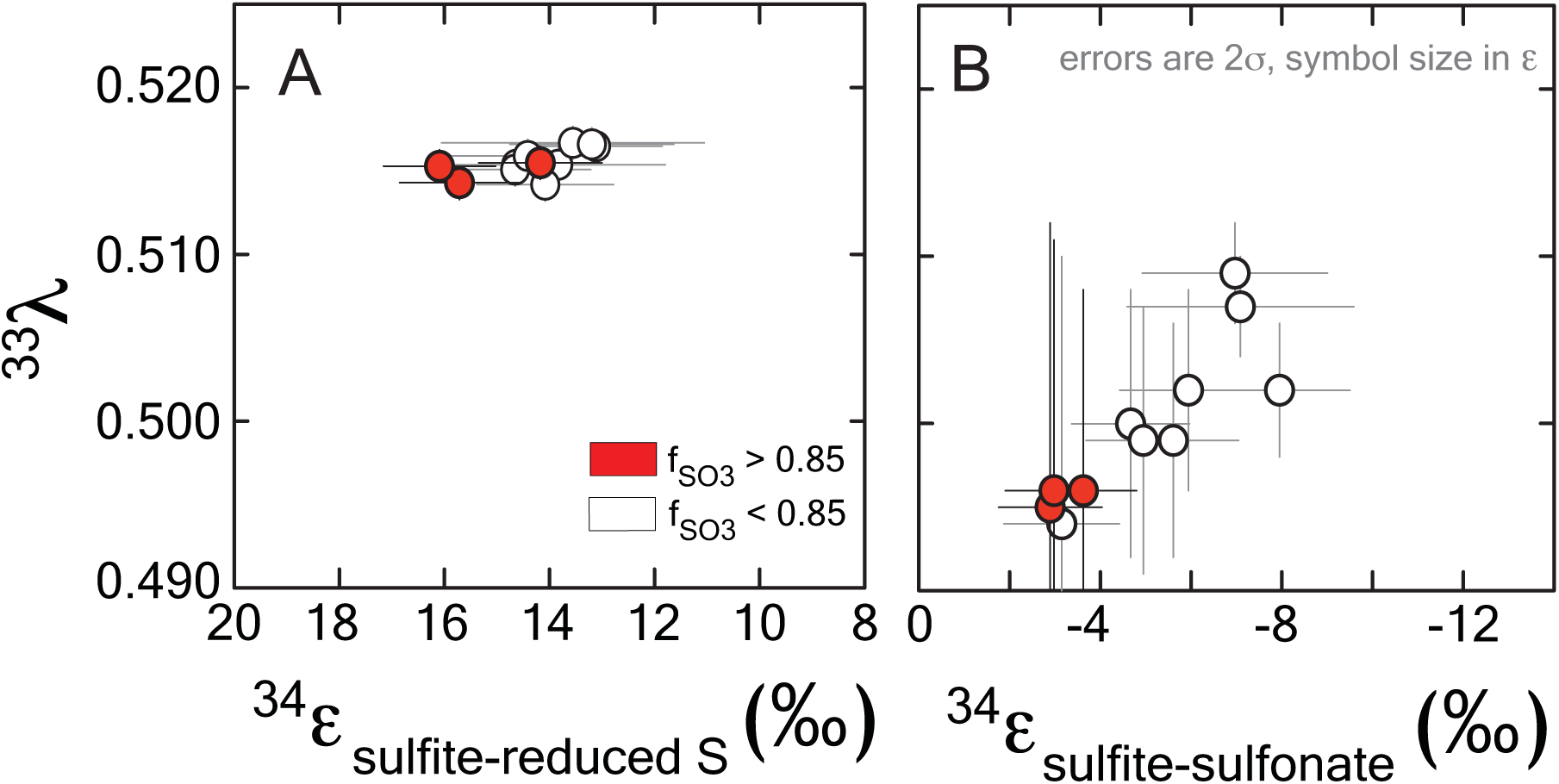
Sulfur isotope fractionation during sulfite reduction by DsrAB. Presented here is the fractionation between sulfite and reduced sulfur (within trithionate and thiosulfate) facilitated by in vivo DsrAB (left frame). Initial values of ^34^ε_DsrAB_ when *f*_SO3_ is > 0.85 are near 15‰. A small amount of variability accompanies changes in *f*_SO3_ to < 0.85. Data from *A. fulgidus* overlap *D. vulgaris* DsrAB data (see Figure S2). The minor sulfur isotope fractionation, ^33^λ_DsrAB_, is stable near 0.515. Fractionation factors for sulfonate generation, presented in the right frame, are much smaller in ^34^ε and carry the opposite sign. Both the ^34^ε and ^33^λ evolve over the time course of the experiment, only after *f*_SO3_ < 0.85. Data and application to calculations are further discussed in the text.

Fractionation of ^33^S between sulfite and reduced sulfur by *D. vulgaris* DsrAB is reported as ^33^λ_DsrAB_, with a calculated result of 0.5150±0.0012 (2σ) over the initial range of *f*_SO3_. The conversion of sulfite to sulfonate yielded a calculated ^33^λ_DsrAB_ that changes as the reaction progressed, from 0.495±0.017 (2σ) at *f*_SO3_ > 0.85, towards 0.510 at *f*_SO3_ < 0.85. The experimental error on ^33^λ_DsrAB_ is inversely related to the magnitude of ^33^ε_DsrAB_ (Johnston et al., 2007), thus is larger for sulfonate generation. We interpret the observed fractionation factors between sulfite and reduced S as representing the binding and reduction of sulfite by DsrAB. The fractionation associated with sulfonate production is more difficult to uniquely diagnose given the wide array of potential biotic and abiotic reactions.

## Discussion

Microbial sulfate reduction is a major process in the sulfur cycle and generates characteristic isotopic fractionations. These fractionations are critical in tracing the movement of sulfur within natural settings (marine and lacustrine). Determining the isotope effects associated with key enzymes in this pathway is critical to disentangling biological and physical controls on the distribution of sulfur isotopes among environmental pools of sulfur. In this study we provide the first constraints (^34^ε and ^33^λ) on the isotope effects associated with one such enzyme: dissimilatory sulfite reductase, the central redox enzyme in dissimilatory sulfate reduction. This experimental constraint provides insight and critical boundary conditions for understanding sulfur isotope fractionation by sulfate reducers. Fortunately, half a century of research on sulfur isotope fractionation by MSR in vivo puts in place a series of useful observations that help to guide our interpretation as to the role of DsrAB. This in turn allows significantly greater access to the information locked in sulfur isotope records.

There is a rich literature of whole cell isotope fractionation data associated with MSR, but information about kinetic isotope effects associated with specific enzymes within the metabolism are entirely lacking. Cellular-level observations include secular and spatial trends in sulfur isotope records attributed to changes in the environmental conditions at the site of MSR, and degree to which biogenic sulfide is preserved in marine sediments (Holland, 1973; Canfield and Farquhar, 2009; Leavitt et al., 2013). The environmental variables most commonly invoked to explain isotopic variability are aqueous sulfate and organic carbon concentrations (Goldhaber and Kaplan, 1975; Habicht et al., 2002; Bradley et al., 2015). Both of these variables ultimately contribute to the net reduction rate and carry independent biological thresholds, one of which ultimately becoming rate-limiting (Bradley et al., 2011; 2015). More specifically, variability in these substrates is manifested as changes in the cell-specific rates of MSR in both the laboratory and natural environment (Goldhaber and Kaplan, 1975; Chambers et al., 1975; Leavitt et al., 2013). In laboratory experiments and natural marine and lacustrine systems, volumetric sulfate reduction rates scale primarily as a function of the availability of sulfate relative to common electron donors like organic carbon (Goldhaber and Kaplan, 1975; Chambers et al., 1975; Leavitt et al., 2013; Sim et al., 2011b). Indeed, sulfate can be non-limiting even in environments with as little as μM sulfate (Nakagawa et al., 2012; Gomes and Hurtgen, 2013; Crowe et al., 2014; Gomes and Hurtgen, 2015; Bradley et al.; 2015), assuming organic matter is more limiting to allow a fractionation to occur (Wing and Halevy, 2014; Bradley et al.; 2015). Constrained whole cell (in vivo) laboratory experiments demonstrate that when electron donors are limiting, the magnitude of fractionation between sulfate and sulfide (^34^ε) carries a nonlinear inverse relationship with cell-specific sulfate reduction rates (Leavitt et al., 2013; Kaplan and Rittenberg, 1964; Chambers et al., 1975; Sim et al., 2011b; Harrison and Thode, 1958). Thus, the range of isotopic compositions produced and preserved in natural environments are interpreted as an output of intracellular rates, which scales with enzyme activity associated with microbial sulfate reduction (Leavitt et al., 2013; Goldhaber and Kaplan, 1975).

In addition to following a rate relationship, fractionation in MSR isotope studies often approaches characteristic upper and lower fractionation limits. Recent experimental work at low sulfate reduction rates captures a ^34^ε_MSR_ (the net isotope effect of microbial sulfate reduction) of nearly 70‰ (Canfield et al., 2010; Sim et al., 2011a). This magnitude of fractionation approaches the theoretical low temperature equilibrium prediction of 71.3 to 67.7‰ between 20° and 30°C (Farquhar et al., 2003; Tudge and Thode, 1950), inspiring research more directly comparing the biologically catalyzed reversibility of MSR enzymes and that of equilibrium (Wing and Halevy, 2014) (see also (Rees, 1973; Holler et al., 2011; Brunner and Bernasconi, 2005; Farquhar et al., 2003; Bradley et al., 2011; Johnston et al., 2007; Farquhar et al., 2008; Mangalo et al., 2008; Bradley et al., 2015)). These studies are fueled by the knowledge that direct (abiotic) equilibration between sulfate and sulfide at Earth surface temperatures is exceedingly slow, with a half-life of exchange estimated at 1.1x10^10^ (at 30°C) to 1.6x10^12^ years (at 20°C; these values are extrapolated from (Ames and Willard, 1951)). Thus, large fractionations between sulfate and sulfide at Earth surface conditions strongly suggests a role for biology.

Most experiments with sulfate reducing microorganisms result in isotope fractionations much smaller than would be predicted from abiotic equilibrium estimates. More than half a century of research and 648 observations from in vivo MSR experiments capture a median isotope fractionation of 16.1‰ (both ^34^ε_MSR_ and in sulfite reduction experiments: Figure 4). In fact, half of experimental data fall between 10 and 22.5‰. This is consistent with the phenomenology of laboratory experiments being conducted at significantly higher sulfate reduction rates than occur in most natural settings. However, given that all these experiments occurred with the same biochemical network, any enzyme-level explanation for the range of fractionations observed at both high and low sulfate reduction rates must be internally consistent.

**Figure 4:**
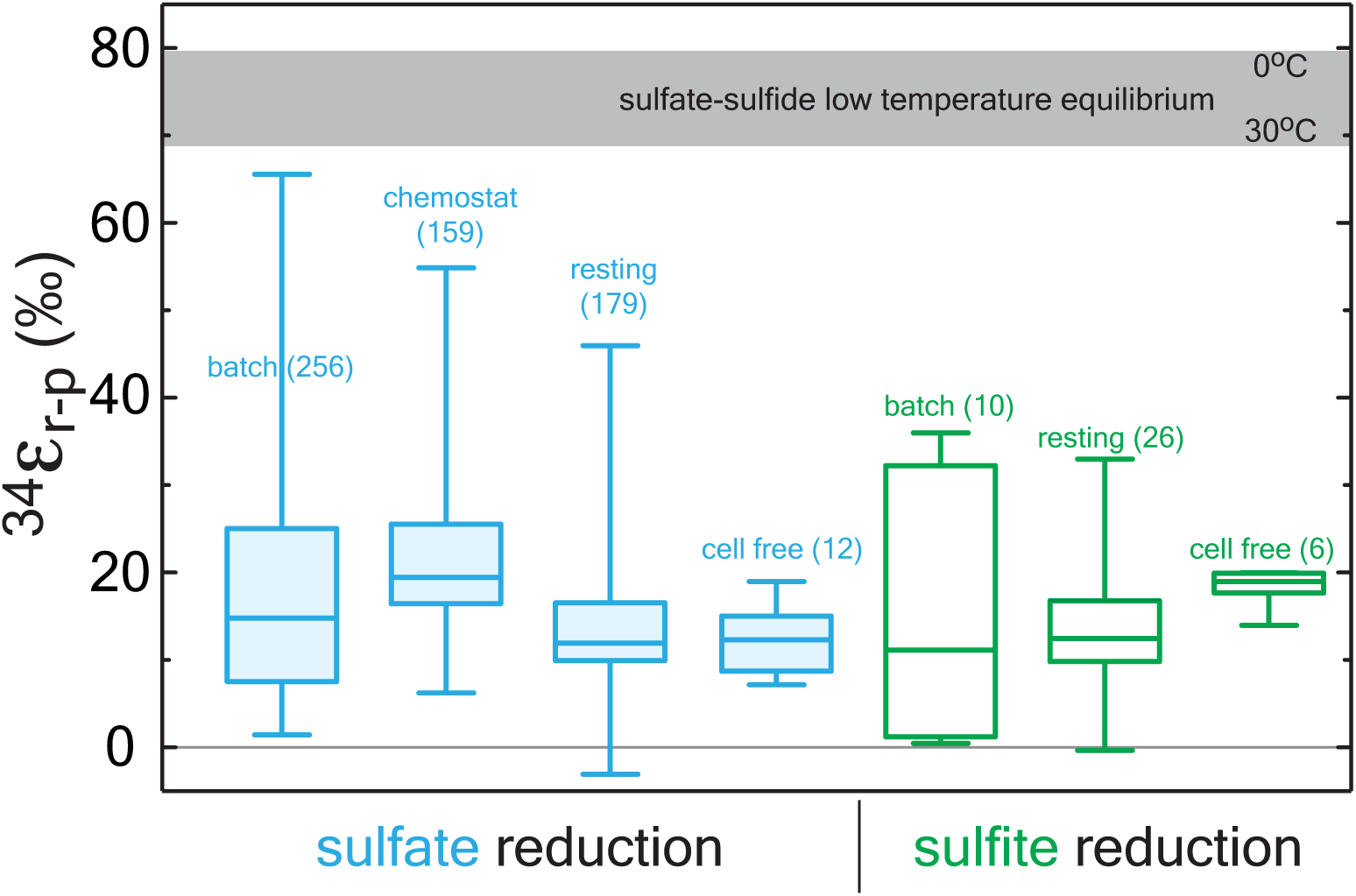
A box and whisker plot of previously published sulfate (blue) and sulfite (green) reduction experiments. All data is binned by experimental approach. The whiskers reflect the entire range of the data, with the boxes reflecting the middle 50% of the data. The median of the data is represented by the bar dividing the box. The bar running across the top is a temperature dependent prediction based on low temperature thermodynamic equilibrium (Farquhar et al 2003). The statistical method and output are detailed in the Appendix along with the compiled data (http://dx.doi.org/10.6084/m9.figshare.1436115), where all compiled values are from the following sources: (Harrison and Thode, 1957; 1958; Krouse et al., 1968; McCready et al., 1975; McCready, 1975; Sim et al., 2012; 2013; 2011b; 2011a; Johnston et al., 2007; Johnston, 2005; Farquhar et al., 2003; Thode et al., 1951; Bolliger et al., 2001; Knöller et al., 2006; Detmers et al., 2001; Jones and Starkey, 1957; Kleikemper et al., 2004; Mangalo et al., 2008; 2007; Canfield, 2006; Hoek et al., 2006; Pallud et al., 2007; Böttcher et al., 1999; Smock et al., 1998; Kemp and Thode, 1968; Ford, 1957; Leavitt et al., 2013; 2014; Chambers et al., 1975; Davidson et al., 2009; Habicht et al., 2005; Kaplan and Rittenberg, 1964).

As described above, a suite of enzymes and cofactors drives dissimilatory sulfate reduction. During the reduction of sulfate to sulfide, sulfur isotope effects are likely to result primarily from transformations that involve the making or breaking S related bonds. Initial steps in sulfate reduction, such as transport into the cell and activation via a reaction with ATP to generate APS (Figure 1 (Fritz, 2002)), do not involve the formation of new S linkages, and are not predicted to be associated with primary isotope effects. Influence on the expressed isotopic fractionation due to transport limitation is, however, conceivable. That is, the concentration of sulfate in the cell may influence the expression of downstream isotope effects, altering the net observed ^34^ε_MSR_. Sulfate transporters may also induce an isotope effect associated with varying membrane fluidity or other strain-specific optima, in response to changing temperature (Kaplan and Rittenberg, 1964; Canfield, 2006), pH (Furusaka, 1961), or as environmental sulfate concentrations become metabolically limiting (Habicht et al., 2005) (see discussion in Bradley et al. 2015).

Primary isotope effects are predicted where bonds are made or broken. APS reductase catalyzes a two-electron exchange that breaks a S-O bond during reduction of APS to generate free sulfite. From the crystal structure of ApsR (Fritz, 2002), it is apparent that the enzyme binds with the APS bound sulfur directly on a nitrogen in the FAD (flavin adenine dinucloetide) cofactor. The product sulfite is then available to interact with DsrAB. This heterodimeric enzyme binds sulfite in an active site containing siroheme. The formation of the Fe-S bond between siroheme and sulfite may be the critical reaction controlling isotope fractionation. Following this, sulfite is proposed to be reduced by the step-wise transfer of two electrons to form S^2+^, then two additional electrons to form S^0^ (Parey et al., 2010). Under in vivo conditions, the S^0^ intermediate was suggested to be withdrawn from the DsrAB complex by the small transfer protein DsrC (Oliveira et al., 2008b; Venceslau et al., 2014). Under in vitro conditions, DsrC is generally absent, and the reduced sulfur in the active site may react with excess sulfite, forming thiosulfate and trithionate (Figure 1) (Drake and Akagi, 1976). DsrC is independently regulated in vivo (Karkhoff-Schweizer et al., 1993), and generates the terminal sulfide from DsrAB bound sulfur derived from sulfite (Venceslau et al., 2014). The relative importance of this protein has only been realized in the last few years (Oliveira et al., 2008b; Venceslau et al., 2014), and has an unconstrained isotope effect.

In general, the magnitude of the thermodynamically predicted sulfur isotope effect scales positively with the number of bonds are made or broken (Tudge and Thode, 1950; Bigeleisen and Wolfsberg, 1958). As described above, sulfite reduction by DsrAB is a central enzyme in MSR, breaking three S-O bonds (Venceslau et al., 2014; Oliveira et al., 2008b), and therefore knowing the fractionation associated with this step is critical to any predictive MSR isotope model (c.f. (Rees, 1973; Brunner and Bernasconi, 2005; Johnston et al., 2007; Farquhar et al., 2007; 2008; Bradley et al., 2011; Wing and Halevy, 2014; Bradley et al., 2015)). Our direct constraint on the fractionation imposed by sulfite reduction indicates that the published assignments of 25‰ (Harrison and Thode, 1958; Rees, 1973) and 53‰ (Brunner and Bernasconi, 2005) for DsrAB are significant over-estimates. It is perhaps not surprising, given that previous appraisals were generated through various indirect approaches (Harrison and Thode, 1958; Rees, 1973; Farquhar et al., 2003; Johnston et al., 2007; Brunner and Bernasconi, 2005), (Harrison and Thode, 1958; Rees, 1973; Farquhar et al., 2003; Johnston et al., 2007; Brunner and Bernasconi, 2005). As previously highlighted, this represents a major limitation to model applications (Chambers and Trudinger, 1979).

Our measured ^34^ε_DsrAB_ value for sulfite reduction (15.3±2.0‰) is large enough to account for the majority of the fractionations observed in the bulk of published the whole-cell MSR experiments over the last sixty-five years (median of 16.1‰, n = 648; Figure 5). As noted previously, laboratory experiments carry a strong bias toward higher rates of sulfate reduction, and as such, the data compilation should be viewed in this light. As most recently articulated through a series of chemostat experiments (Leavitt et al., 2013; Sim et al., 2011a), the consequence of elevated metabolic rate is a smaller relative ^34^ε. In isotope biogeochemistry, relationships like this often depend on the single slowest overall rate-limiting step within a metabolism (Mariotti et al., 1981; Hayes, 1993). The fractionation limit at high metabolic rates in cultures (^34^ε = 17.3±1.3‰), marine sediments (^34^ε = 17.3±3.8‰) and DsrAB are statistically indistinguishable (Figure 5). This similarity is consistent with DsrAB as a rate-limiting step explaining the majority of observed fractionation (Figure 5). However, this interpretation omits complexity associated with the metabolic network.

**Figure 5:**
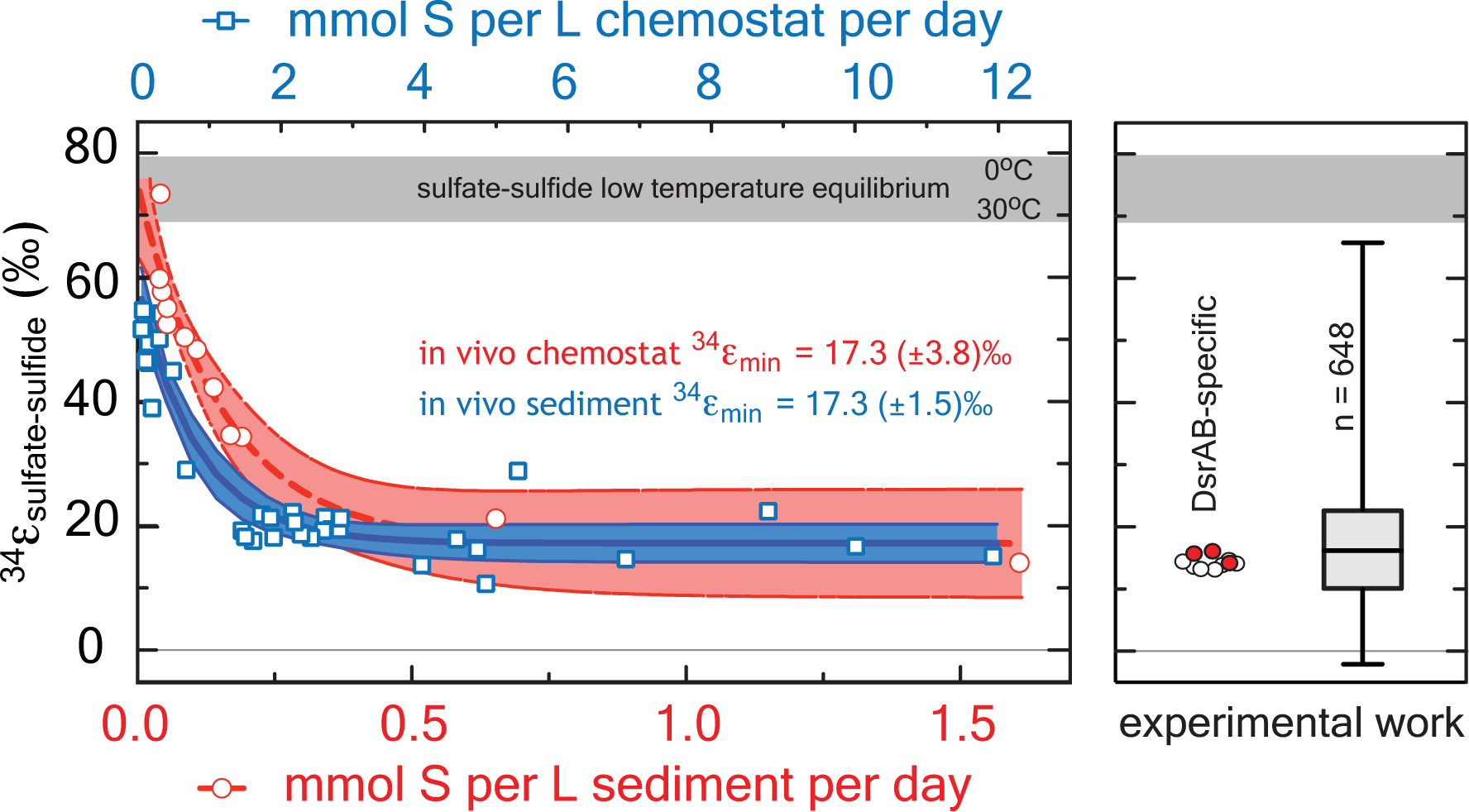
A comparison between modern marine (environmental) and laboratory (experimental) S isotope fractionations, as a function of sulfate reduction rate. These data are further referenced to a statistical distribution of published experimental fractionation data. (A) Fractionation as a function of volumetric sulfate reduction rate from axenic continuous culture experiments (blue squares and regression) (Leavitt et al., 2013) and modern marine sediments (red circles and regression) (Goldhaber and Kaplan, 1975). Solid lines are mean values with shaded regions representing the 95% confidence interval around a non-linear regression. While the upper fractionation limits are offset, perhaps due to differences in biomass per volume of sediment versus volume of chemostat, the limits approached at high reduction rates are statistically indistinguishable at 17.3‰. (B) The fractionation associated with DsrAB experiments, color-coded as in Figure 3 and on the same isotope scale. Also included is a box-whisker treatment of all measured ^34^ε (n = 648) sulfate and sulfite reduction experiments compiled in Figure 2. Here, the median value is 16.1‰, also statistically indistinguishable from chemostat and modern marine sediments limits at elevated rates of sulfate reduction. Included for reference is the theoretical sulfate-sulfide equilibrium fractionation (gray bar) for 0 to 30°C (Farquhar et al., 2003).

This raises an essential question: how does the DsrAB constraint change our understanding of the possible range of fractionations imposed by MSR? For instance, if we assume that DsrAB is the slowest reaction in MSR, then the metabolic steps preceding sulfite (SO_4_^2-^ ⇔APS ⇔ SO_3_^2-^) will necessarily approach equilibrium (Wing and Halevy, 2014). The thermodynamic predictions would then require an accompanying fractionation approaching 25‰ between sulfate and sulfite at biologically relevant temperatures (the equilibrium fractionation estimate (Farquhar et al., 2003)). At its simplest, this effect would be additive with that of DsrAB (25 + 15.3 ‰, or 40.3‰), which encompasses a majority of experimental MSR isotopic fractionations. However, the lower plateau in fractionation approached at high rates, stemming from the calculations using both modern marine sediment and chemostat data (see the Appendix) – is less than half of this magnitude. That is, the lower plateaus for in vivo fractionation at high rates of 17.3‰ allows only a ∼2‰ fractionation partitioned among these upstream steps, if in fact DsrAB is fully expressed. This smaller ‘upstream’ kinetic fractionation is, however, consistent with the few loose estimates from crude cell extracts and resting cell studies (i.e. not purified enzymes), which putatively suggests a ^34^ε of 4 to 15‰ for the cumulative activation of sulfate to APS and reduction to sulfite (Ford, 1957; Kemp and Thode, 1968). This may suggest that the sulfate-sulfite conversion (in vivo and in vitro) reflects a predominately kinetic (rather than equilibrium) control. This requires, however, we loosen the degree to which DsrAB is called upon to fully control the net MSR isotopic fractionation, invoking some delicate balance between upstream reactions and that of DsrAB, but maintaining the model where enzyme kinetics (especially DsrAB) will win out over equilibrium effects as sulfate reduction rates move from low to high.

An alternate explanation is that APS reductase (ApsR) is the rate-limiting step under sulfate replete conditions (Rees, 1973). If this is the case, fractionation imposed by DsrAB is unexpressed, as it is downstream of ApsR (Hayes, 2001; Rees, 1973). If fractionation associated with APS reductase is near 17‰, it could alone account for most of the observed fractionation observed at high sulfate reduction rates. The fractionation imposed by ApsR has not been directly measured, though can be proxied from the discussion above. However, evidence against ApsR as the rate-limiting step is shown by studies indicating reversibility of the ApsR (Peck, 1960) and the sulfate reduction pathway (Chambers and Trudinger, 1975; Holler et al., 2011). Recent studies using oxygen isotopes as tracers have demonstrated that some intracellular sulfite is oxidized in vivo back to sulfate (Mangalo et al., 2007; Farquhar et al., 2008; Turchyn et al., 2010; Einsiedl, 2008; Mangalo et al., 2008). These studies demonstrate that sulfite re-oxidation is commonplace in MSR and often quantitatively significant (Antler et al., 2013). This reoxidation is inconsistent with ApsR being rate-limiting under the range conditions tested.

Any explanation for the net MSR isotopic fractionation must also account for the large fractionations observed at low sulfate reduction rates. These large fractionations are common in nature, and require another type of mechanism. These isotopic fractionations approach but do not reach the theoretical equilibrium values for sulfur isotope exchange between sulfate and sulfide (Figures 4 and 5) (Farquhar et al., 2003; Tudge and Thode, 1950; Johnston et al., 2007). In this context, further analysis to understand intracellular thermodynamics is critical (the redox pairs responsible for various reactions, see (Wing and Halevy, 2014)), along with measurements of the intrinsic isotope effects of other key enzymes in the metabolic network, including ApsR and DsrC. In that sense, this study represents a key first step.

In parallel to examining the ^34^ε effects, measuring minor S isotope (^33^S/^32^S) fractionation provides additional information about the class of reaction mechanism associated with in vitro DsrAB activity. In our experiments, the conversion of sulfite to sulfonate carries a ^33^λ of from ∼ 0.496 (±0.012, 2σ), evolving toward 0.510 as the reaction proceeds. Sulfite reduction via DsrAB has an invariant ^33^λ of 0.5150 (±0.0012, 2σ). In cases where ^33^λ is 0.515, a purely equilibrium fractionation is often inferred but not required, while values less than this require kinetic effects (Young et al., 2002; Farquhar et al., 2003). In this framework, in vitro sulfonate formation falls under kinetic control, while formation of reduced S could be interpreted as a kinetic or equilibrium reaction. Thus, specific predictions for the DsrAB enzyme require more detailed modeling of the structure and function of the DsrAB enzymatic active site. This level of analysis – where inroads joining empirical work with theory are constructed – is present in analogous systems (Karsh et al., 2012), but absent within the S cycle. Nonetheless, the work here provides the only triple-isotope constraints on enzyme-specific fractionation factors in both MSR and the global biogeochemical S cycle. Further, this approach may prove useful in other enzymatic systems where elements with ≥ 3 stable isotopes are involved (e.g. O, Fe, Ca, Mg, Se, Zn, Mo).

## Conclusions

Direct constraints on enzymatic isotope effects, when placed in context of laboratory and field observations, represent a key step towards improving our understanding of how environmental factors come to control biochemical sulfur isotope fractionations in nature. Experimental results indicate that the kinetic isotope effect generated by dissimilatory sulfite reductase, the enzymatic core of MSR, generates less than a quarter of the maximum fractionation observed in sulfate reduction experiments and modern marine sediments. However, the ^34^ε_DsrAB_ aligns nicely with the vast majority of experimental data generated over the last 65 years, as well as chemostat and marine sediment studies sampling high rates of sulfate reduction. The consistency between these published fractionations and the DsrAB isotope effect suggests a fundamental role of this enzyme in setting sulfur isotope compositions. This work highlights the need for further consideration of the allied enzymes in MSR and the likelihood of abiological (and/or equilibrium) effects as microbial sulfate reduction rates slow. Though questions remain, placing quantitative constraints on the core of sulfate reduction – DsrAB – represent a fundamentally new direction in exploring experimental and environmental sulfur isotope records today and throughout Earth history.

## Acknowledgments

Thanks to Sofia Venceslau and Fabian Grein for discussions on MSR biochemistry and DsrC cycling; Isabel Pacheco for the purified hydrogenase; to Erin Beirne and Andy Masterson for expert assistance in measurements and separation chemistry. Thanks to Ann Pearson, David Fike and Itay Halevy for comments that greatly improved our interpretations and presentation. We especially thank John Hayes for exceptionally in-depth feedback. This work was funded by an NSF-GRFP (WDL), NSF Geobiology and Low-Temperature Geochemistry (DTJ, IACP), the Sloan Foundation (DTJ), the Agouron Institute (ASB), PTDC/QUI-BIQ/100591/2008 and PTDC/BBB-BQB/0684/2012 (IACP), UID/Multi/04551/2013 (to ITQB) funded by Fundacao para a Ciencia e Tecnologia (FCT, Portugal), Washington University in St. Louis (ASB) and the Steve Fossett Postdoctoral Fellowship at Washington University in St. Louis (WDL).

## APPENDIX

### Appendix 1 Operational definitions of S moieties

In this study we measured the concentrations of three pools: sulfite, trithionate, and thiosulfate; hydrogen sulfide was not detected. We measured the major and minor sulfur isotopic compositions of three operationally defined pools: ‘reactant’ sulfite (initial and residual), product ‘sulfonate’, and ‘reduced product’ S. What we refer to as the pooled product ‘sulfonate’ sulfurs are known in inorganic chemistry as sulfuryl groups (O_2_S-X_2_), where one of the X’s represents an O^-^/OH and the other a S in oxidation state 0 (trithionate) or -1 (thiosulfate), meaning the outer sulfuryl-S’s are in approximately oxidation state +5 (thiosulfate) or +4 (trithionate), with initial and residual reactant sulfite sulfur in the standard +4. The sulfonate S differs from sulfite S in that it is bound to either an approximately -1 valent sulfur in thiosulfate (S-S(O)_3_^2-^) (Vairavamurthy et al., 1993) or as two sulfonates each bound to one sulfur of valence approximately 2+, in trithionate ((O_3_S-S-SO_3_)^2-^). In this study we refer to the 0 and -1 oxidation state sulfurs from trithionate and thiosulfate, respectively, as the ‘reduced product’ S pool. They are grouped by our operational extraction (see below). For explicit definitions and nomenclature refer to the ‘IUPAC Goldbook’ (McNaught and Wilkinson, 1997).

### Appendix 2. Enzyme purification and in vitro experiments

#### A2.1 DsrAB isolation and purification

DsrAB was purified from *Desulfovibrio vulgaris* Hildenborough (DSM 644) and *Archaeoglobus fulgidus* cells grown in a 30 or 300L batch culture in a modified lactate/sulfate medium (Oliveira et al., 2008a) at iBET (Instituto de Biologia Experimental Tecnológica; www.ibet.pt), grown at 37 or 80°C, respectively. The soluble cell fraction was obtained as previously described (Le Gall et al., 1994; Oliveira et al., 2008a). All purification procedures were performed under atmosphere at 4°C using an AKTA FPLC (Amersham Biotech Pharmacia) with two buffers, (A) 20mM TrisHCl and (B) 50mM TrisHCl with 1M of NaCl (both pH 7.6 and containing 10% glycerol). Buffer (A) was used to equilibrate the columns and buffer (B) to generate the ionic strength gradient. The soluble cell fraction was loaded into a Q-Sepharose fast-flow (XK50/30) column, and a stepwise salt gradient applied, with the DsrAB-containing fraction eluting at 300 mM NaCl. The characteristic DsrAB (‘desulfoviridin’) absorption peak at 630 nm was used to track the protein, as previously described (Marritt and Hagen, 1996; Wolfe et al., 1994). DsrAB-containing fractions were then loaded into a Q-Sepharose fast-flow (26/10) column and eluted in 250 mM NaCl. To verify enzyme purity, the final DsrAB-containing sample was analyzed by 12% SDS-PAGE gel electrophoresis. DsrC is present in the DsrAB preparation from *D. vulgaris*, but remains functionally inactive during in vitro assays as previously described (Oliveira et al., 2008b), and also due to the lack of DsrMKJOP (Venceslau et al., 2014). Thus, we refer only to the ‘DsrAB’ fraction in the *D. vulgaris* experiments. In the *A. fulgidus* experiments, DsrAB is free of any DsrC. Protein was quantified by the method of Bradford (Bradford, 1976). The *Desulfovirio gigas* [NiFe] hydrogenase used in all assays was purified as described previously (Romão et al., 1997).

To ensure the activity of purified DsrAB was not strongly influenced by the high initial concentration of sulfite used in the fractionation experiments (10 or 15mM), we performed small-volume kinetic assays under the same conditions as for isotope measurements. Sulfite alone was measured by HPLC on monobromobimane (MBBr) derivatized samples (Newton et al., 1981). Once the sulfite concentrations for each initial and final (0 and 2 hours) time points were sampled, derivatized, measured and calculated, we applied a non-linear regression formulated from the standard Michaelis-Menten equation, solving for the V_max_ and K_m_.

#### A2.2 *D. vulgaris* DsrAB in vitro fractionation experiments in detail

To determine the DsrAB-specific S isotope fractionation factors we designed and executed a series of batch (closed-system) sulfite reduction experiments. The key considerations in experimental design are: (*i*) to provide enough sulfite at t_0_ to ensure we generate significant enough quantities of all the product pools so we can measure, at high precision and accuracy, and at multiple [time] points on *f*, the multiple S isotopic composition of each pool (*i.e.*, 2μmols of S per pool per SF_6_ measurement on the DI-IRMS, which means 2μmols per fluorination reaction); (*ii*) to provide the proper reaction conditions to allow DsrAB optimal activity for goal *i* (pH = 7.1, T = 20 or 31**°**C); (*iii*) to ensure hydrogenase activity is not inhibited by the pH chosen (optimum above pH 7.5, activity significantly depleted below pH 6.5, so we chose pH 7.1, to account for optima of both DsrAB and [NiFe]-Hydrogenase); and finally (i*v*) to ensure the sulfite to hydrogen ratio favors sulfite over reductant capacity (i.e., *p*H_2_ in the headspace relative to [sulfite]_t0_), such that no more than 75% of t_0_ sulfite is consumed to all products, and less than 50% to the reduced S (dictated by the amount of H_2_ in the headspace. Finally, (*v*) determining the sampling interval to ensure proper distribution of points along *f*, such that applying a closed system distillation model is possible, and statistically robust. Data plotted in Figure 3 represents experimental results that met all of these conditions. The full experimental results (33 experiments) are contained as a supplemental file (http://dx.doi.org/10.6084/m9.figshare.1436115).

All experiments were prepared in an anaerobic chamber. In vitro reactions were carried out in 100mL acid-washed, autoclave-sterilized, borosilicate glass bottles sealed with butyl-rubber septa and aluminum crimps. Each bottle contained 50% reaction buffer and 50% gaseous headspace. This was done to sufficient H_2_ was present in the headspace to reduce at most 50% of the sulfite, based on an estimate using Henry’s law and available solubility constants for H_2_ at the given preparation temperatures and headspace pressures. During manipulations in the anaerobic chamber the chamber gas was initially 95:5 N_2_:H_2_. Upon removing the reaction mixture-filled bottles from the chamber, these were capped and crimped, and headspace completely exchanged with deoxygenated 100% Ar, then finally exchanged for 100% H_2_ to initiate the experiments. Experimental buffer is 50mM phosphate buffer (KP*i*) prepared at pH 6.9±0.05, with final pH is 7.1±0.05 following the addition of the stock Na_2_SO_3_ solution (the reaction is therefore initiated at 7.1). All reaction solutions contained the following: 50mM KP*i* buffer (final pH ±0.05), 10 or 15 mM sodium sulfite, 0.832mM methyl viologen, 242 nM or 315 nM of *D. vulgaris* DsrAB (calculated to give the same activity depending on the DsrAB aliquot selected), and 8.25 nM [NiFe] hydrogenase (297 U/mg). All experimental mixtures and reagents were prepared in previously boiled 18.2MΩ water, cooled under O_2_-free N_2_.

#### A2.3 *A. fulgidus* DsrAB in vitro fractionation experiments in detail

To extend our studies to a different taxonomic form of the enzyme, we used DsrAB from the thermophilic archaeon *A. fulgidus.* This enzyme operates at higher temperature and does not have DsrC present in the complex (Schiffer et al., 2008). The results from these experiments are significantly limited compared to those with *D. vulgaris*, due to too few time points to apply the closed-system model (specifically due to significantly low sample sizes of reduced S for isotope measurements, note the effort made to correct the two data points on *A. fulgidus* reduced-S). Nevertheless, the results obtained are comparable, when considering the measured δ^34^S. The values for these experiments are presented with *D. vulgaris* values in Figure A2.

*A. fulgidus* DsrAB experiments were conducted 15mM initial sulfite and 65°C, and showed consistent loss of sulfite and accumulation of products between replicates. We selectively precipitated, separated, and directly measured the ^32^S-^33^S-^34^S-^36^S compositions from the residual reactant (‘sulfite S’) and the ‘sulfonate S’ ((SO_3_)_*x*_). Only the ^34^S/^32^S compositions of the ‘reduced product S’ [(S)_*y*_] reservoirs were measured (again, due to significantly small reduced-S samples recovered). From these experiments we were able to get a complete set of samples (i.e. sulfite, sulfonate and reduced S) from one time-point and partial sets from another (i.e. sulfite and sulfonate). We are unable to calculate the ^34^ε_DsrAB_ for *A. fulgidus* directly using our modelof sulfite reduction experiments due to the dearth of time points (points on *f*). However, the *A. fulgidus* isotope values agree with those measured sulfite, sulfonate and reduced S moieties for *D. vulgaris* (Figure A2). This general agreement between *D. vulgaris* and *A. fulgidus* DsrAB, independent of temperature or phylogenetic origin is perhaps unsurprising, given that previous theoretical predictions deemphasize the role of temperature in determining the magnitude of *kinetic* isotope effects (Bigeleisen and Wolfsberg, 1958). Furthermore, these enzymes tightly share active site structures (Oliveira et al., 2008b; Parey et al., 2013).

### Appendix 3. Analytical methods & data handling

#### A3.1 Quantification of dissolved species

To quantify sulfite and bisulfite concentration in solution we adapted a protocol to quantify SO_2_ dissolved in water (Grant, 1947), referred to as the ‘Fuschin’ assay from here foreword. Our protocol is specific to the in vitro DsrAB assay conditions. It was determined that matrix matching between samples and standards and the exclusion of oxygen is critical to a successful and reliable assay. Furthermore, we determined trithionate, thiosulfate, sulfate, and zinc sulfide solids do not interact with this color-reagent in the assay. The Fuschin assay is useful over a range of 0-40 nanomoles of sulfite in the final assay volume of 1mL. Standards of sodium sulfite (Na_2_SO_3_ anhydrous, analytical grade) were prepared immediately before the assay is performed in deoxygenated water (boiled and degassed with N_2_) or Kp*i* buffer. The reaction mixture is composed of 0.04% w/v Pararosaniline HCl (analytical grade) in 10% H_2_SO_4_ (analytical grade) v/v, prepared stored in an aluminum-foil wrapped tube or amber-glass bottle at 4°C; and 3.7% formaldehyde (HCHO) prepared fresh each day by diluting 37% (stock) formaldehyde 1:10 water. The reaction is performed on the bench working under N_2_ flow, or in an anaerobic chamber. A detailed step-by-step protocol is available in (Leavitt, 2014).

Trithonate and thiosulfate were measured by a modified cyanolysis protocol (Kelly and Wood, 1994; Kelly et al., 1969; Sörbo, 1957). We primarily employed the method of Kelly and Wood (Kelly and Wood, 1994) modified in the following manner: the reaction volumes were reduced to 10 rather than 25 mL’s (still in volumetric flasks) and we used nitric rather than perchloric acid. Nitric acid was used in the original version of this method (Sörbo, 1957), allowing us to avoid the significant hazards of working with significant volumes of perchloric acid. Samples were added to the reaction buffer to fit within the range of ferric thiocyanate standards (prepared from potassium thiocyanate as a simple standard and thiosulfate as a reaction standard) from 5 to 25 μM (final concentration in the 10mL reaction), as well as ‘blanks’ prepared from the in vitro assay reaction buffer (50 mM potassium phosphate buffer at pH 7). This is typically 400 μL of in vitro solution added to the 10 mL cyanolysis reaction, in duplicate per method (2X thiosulfate determinations and 2X trithionate determinations). A detailed step-by-step protocol is available in (Leavitt, 2014).

For sulfide quantifications, we preserved samples in zinc acetate (2% w/v) from each closed system reaction, using a modified Cline (Cline, 1969) method. Analytical grade sodium sulfide (>98.9% Na_2_S*9H_2_O) was used as the standard, and prepared in deoxygenated (boiled and N_2_-sparged) in vitro reaction buffer, by precipitating the sulfide with excess zinc acetate (anhydrous), mimicking our sampling protocol. A detailed step-by-step protocol is available in (Leavitt, 2014). In all samples no sulfide was detected above the determined detection limit of 6.25 μM. From the literature reports where membrane fractions were omitted (Drake and Akagi, 1978), signifying a lack of DsrC re-cycling mechanism (Oliveira et al., 2008b; Bradley et al., 2011), we expected little to no sulfide. This is further supported by the closure of S mass balance at each time-point from each experiment, within analytical error (see main text). Blanks were prepared identically to those in the cyanolysis protocol.

#### A3.2 Sample preparation for S isotope analysis

All S-bearing samples for S-isotope analyses (ultimately as SF_6_ and/or SO_2_) were removed from the in vitro reaction solution (50mM potassium phosphate) following a sequential precipitation protocol, inspired by that of Smock and colleagues (Smock et al., 1998). Our protocol reflects our specific experimental setup and the pools we aimed to isolate and purify: sulfite (and any trace sulfate), sulfonate, and reduced product S. Samples of the residual reactant (sulfite) and pooled products (reduced product S and sulfonate S) were removed from the in vitro reaction mixture by sequential precipitation and filtration or centrifugation to isolate solid-phases Ag_2_S_(s)_ (reduced product), BaSO_3(s)_ (sulfite), BaSO_4(s)_ (sulfonate) by the extraction scheme modified from our recent work (Leavitt et al., 2014) and detailed elsewhere (Leavitt, 2014). Samples were captured as BaSO_3(s)_, BaSO_4(s)_ or Ag_2_S_(s)_, respectively. Sub-samples of the reduced product Ag_2_S were directly fluorinated (after the below washing steps were carried out to ensure clean Ag_2_S), or in the case of the sulfonate S-pool, collected from the AVS residue and converted from BaSO_4_ to Ag_2_S by the method of Thode (Thode et al., 1961), according to the protocol we recently published (Leavitt et al., 2013). Novel to this study: all sulfite samples (reacted as BaSO_3(s)_) were oxidized with peroxide prior to ‘Thode’-reduction (detailed protocols for these methods are available in (Leavitt, 2014)). All samples entering the elemental analyzer isotope ratio mass spectrometer (EA-IRMS), and combusted to and analyzed as SO_2_, are prepared as dry BaSO_3(s)_, BaSO_4(s)_ or Ag_2_S_(s)_. All samples for quadruple S isotope analysis enter the fluorination line as pure dry Ag_2_S_(s)_.

#### A3.3 Major S-isotope (^34^S/^32^S) ratios measurements

<u>C</u>ontinuous <u>f</u>low <u>i</u>sotope <u>r</u>atio <u>m</u>ass <u>s</u>pectrometric (CF-IRMS) measurements of the three S-bearing pools of interest, sulfite, sulfonate and reduced product S, were performed as follows: 0.4mg (±0.05mg) BaSO_3_, BaSO_4_ or Ag_2_S were converted to SO_2_ by combustion at 1040°C in the presence of excess V_2_O_5_ (Elemental Analyzer, Costech ECS 4010) and analyzed by continuous flow isotope ratio mass spectrometry (SD = ±0.3‰; Thermo-Finnegan DELTA V Plus). All samples yielded clean chromatography and most m/z 66 amplitudes (corresponding primarily to the (^16^O^34^S^16^O)^+^ ionization product) within the range of in-run standards (IAEA: S1, S2, and S3 for Ag_2_S or SO5, SO6, and NBS-127 for BaSO_4_ and BaSO_3_). Some sulfite precipitates (BaSO_3_sulfite_), though not any of the sulfate or sulfide precipitates (BaSO_4_sulfonate_ or Ag_2_S__reduced product_) produced atypical weight to m/z 66 response to what we regularly note with lab standards of BaSO_3_ or BaSO_4_ – specifically the signal was less than predicted, likely due to occlusion of phosphates (from the in vitro reaction buffer) in the barium sulfite matrix. As a result, we use the m/z 66 to BaSO_3_ weight ratio (mg of BaSO_3_ per unit area of the m/z 66 peak) to calculate the desired sample weight to achieve standard signal size, re-weighed and re-combusted/measured the requisite samples, and in all cases achieved m/z 66 peak areas in the range of our IAEA BaSO_4_ and in-house BaSO_3_ standards. Each standard is measured at least 4x in-run and each sample 2-3x (when sufficient sample is available). This simplifies the scale-conversion calculation for taking samples referenced in-run to in-house standard tank gas (HAR1_SO2_) and ultimately to the international reference frame (V-CDT).

#### A3.4 Multiple S-isotope (^33^S/^32^S, ^34^S/^32^S, ^36^S/^32^S) ratio measurements

Duel-inlet (DI-IRMS) measurements of all four stable S isotopes (^32^S, ^33^S, ^34^S, ^36^S) from the three S-bearing pools of interest, sulfite, sulfonate and reduced product S, were performed as previously described (Leavitt et al., 2013; 2014). Briefly, all samples for quadruple S-isotope analysis, prepared dry and clean Ag_2_S (described above), were fluorinated under 10X excess F_2_ to produce SF_6_, which is then purified cryogenically (distilled at -107°C) and chromatographically (on a 6’ molecular sieve 5Å inline with a 6’ HayeSepQ 1/8”-stainless steel column, detected by TCD). Purified SF_6_ was measured as SF_5_^+^ (*m/z* of 127, 128, 129, and 131) on a Thermo-Finnegan Scientific MAT 253 (SD: δ^34^S ±0.2, δ^33^S ±0.006‰, δ ^36^S ±0.15‰). All isotope ratios are reported in parts per thousand (‰) as experimentally paired sulfates and sulfides measured. Long-term running averages and standard deviations are calculated from measures of IAEA standards: S1, S2, S3 for sulfides or NBS-127, SO5, SO6 for sulfates. Isotope calculations and notation are detailed in the text. Standard deviations for each value is estimated as reported previously (Johnston et al., 2007) with previous inaccuracies in the transcription corrected here.

#### A3.5 Scale-compression correction calculations for small S samples

Experiments with *A. fulgidus* yielded small amounts of product (0.35 to 0.01 mg), which required additional data handling during isotope analysis. Given the small size of these samples, we ran each sample only once and bracketed the samples (n = 2) with a series of standards: IAEA S1, S2 and S3 (n = 16, 14, and 14, respectively) run over a size series that captured the sample sizes, all by CF-IRMS only. As expected, we observed that the measured isotopic composition of the standards varied non-linearly as a function of signal intensity (monitored as peak integrated areas and peak intensities on m/z 64 and 66 for SO_2_^+^ and 48 and 50 for SO^+^). The size dependence on the isotopic composition (handled as ^50^R and ^66^R, which are the 50/48 and 66/64, respectively) scale compression is calculated as a proportional change. For SO (*correction factor* in Figure A3) it scales as: 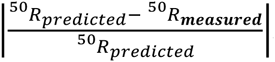. We focus on SO here as these samples yielded sharper chromatography on slightly different sized signals (due to resistor differences between SO and SO_2_ cups – 3 x 10^10^ Ω and 1 x 10^10^ Ω respectively). Thus, using the three IAEA standards, we developed a correction whereby we solve (in the standards) for the non-linear features of the data as it relates to signal intensity (here monitored as the peak integrated area on mass 48 – ^32^SO). This is shown in Figure A3. After this correction is applied to ^50^R_measured_, SO data is converted to an SO_2_ scale (see Figure A3), which is a linear transfer function again derived from IAEA standard data. The final correction places all the data (now on a SO_2_ scale against in-house reference SO_2_ tank gas) to the V-CDT scale. To review, we perform the following steps 1) correcting the ^50^R on SO for sample size, 2) convert ^50^R to ^64^R (against tank gas), and finally 3) convert all data to a VCDT scale.

The largest source of error in this treatment is associated with the sample size correction. As such, we propagate the error associated with the fit in Figure A3 to determine the uncertainty in the final isotope value. As expected, for small samples this error is quite large (Figure A2), with the value decreasing in absolute magnitude as signal intensity (peak integrated area) increases. We also compare these error estimates to the calculated shot noise for this measurement (pink line in Figure A3d). As is presented below, our regressed error is in excess of the shot noise limit. Similarly, the error on the population of standards that were used in deriving this fit is ∼1‰ (n = 44).

### Appendix 4.

#### A4.1 The closed-system distillation model for a more complex network

There exists the possible mixing of multiple fractionation factors later in the experiment (f < 0.85). The approach outlined in the main text yielded results in which the observed fractionation factor between sulfite and reduced pools appeared to change as a function of *f*, when f < 0.85 – that is, later in the reaction, when back-reactions are more likely (Figure A4). One explanation for this apparent behavior is that the reduced pool is not the product of a single set of reactions but of multiple reactions. The most plausible explanation for this is that some fraction of the reduced pool is derived from the sulfonate pool rather than being derived solely from sulfite, particularly later in the in vitro experiment (*i.e.* at values of *f* < 0.85). Previous work (Parey et al., 2010; Drake and Akagi, 1978; 1977) has demonstrated that DsrAB is capable of reducing trithionate to thiosulfate, and thiosulfate to sulfite and sulfide, which was confirmed with the *D. vulgaris* enzyme.

We can constrain the magnitudes of the fractionation factor related to the conversion of the sulfonate to reduced S through the following steps. First, utilizing the framework given above to solve for α_red_ for the time points where *f* is nearest to 1. As these measurements are obtained at the lowest concentrations of product, we assume that this result gives an estimate for α_red_ that reflects the production of reduced S from sulfite only, with minimal input of reduced S derived from sulfonate. Second, we write an equation for *R*_*red*_ as a function of α_red_, *R*_SO3_, and *R*_*ox*_:

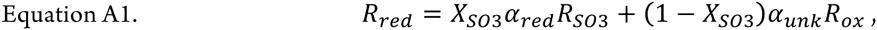

where X_SO3_ is the fraction of *R*_red_ generated directly from sulfite and α_unk_ is the unknown fractionation factor between *R*_ox_ and *R*_red_. This equation is then rewritten and solved for α_unk_ as a function of the other parameters over a range of values of X_SO3_ (0.01 – 0.99). This does not yield a unique solution for the unknown fractionation, but constrains its value given the relative importance of the contribution to the reduced sulfur pool of both sulfonate and sulfite. We assume that the *relative* contribution of the secondary reaction is invariant over the course of the reaction, thereby manifesting as no change in *j*.

#### A4.2 Error propagation calculation for the closed system model estimates

The error associated with calculations of ^33^λ (approximately the slope of a δ^33^S vs. δ^34^S line) is highly sensitive to the length of the line (total ^34^S range, ^34^ε) and modestly related to the residual around a mass-dependent theoretical prediction (the standard deviation on δ^33^S is often used here (Johnston et al., 2007; Farquhar et al., 2003). To approximate the standard deviation (σ) associated with our ^33^λ calculation, we propagate our measurement errors (δ^34^S, concentration, etc.). We keep with the presumption that mass-dependence will dictate the δ^33^S, once the new δ^34^S is calculated. This stems from the fact that the error in δ^34^S and δ^33^S are highly correlated, meaning that the error in δ^33^S is significantly smaller (∼0.008‰) than that for δ^33^S (∼ 0.1‰). As our fractionation factor model is based on a closed-system distillation equation (see above), we perform an error propagation on an equation of the form: R_f_= (R_0_) (*f* ^(α-1)^), where we are most interested in accounting for the analytical errors on the isotope measurement (σ_R_, 0.2‰/1000) and the uncertainty on *f*. The second term is critical here as we are independently determining *f* from concentration measurements in the experiment, with a standard deviation on sulfite concentration measurements of 3%. We use this value moving forward as a metric of σ_f_. To simplify the presentation, we let X = (α -1) and Z = *f*^X^. Following typical error propagation for power law and multiplicative relations (Bevington and Robinson (2003) page 43-46) (Bevington and Robinson, 2003), we find:

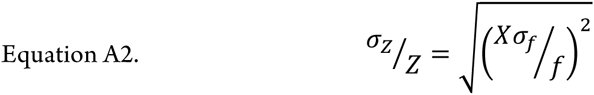

which then can substitute into the final form of:

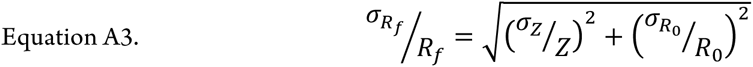

The σ_Rf_ is then converted into ‰ units (through multiplying by 1000) so that it can be inserted into the updated (Johnston et al., 2007) error equation for ^33^λ, presented here:

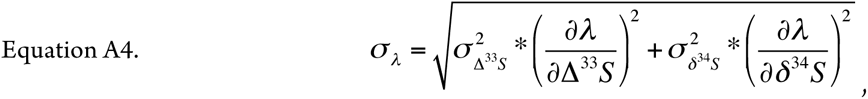

which can be broken down into:

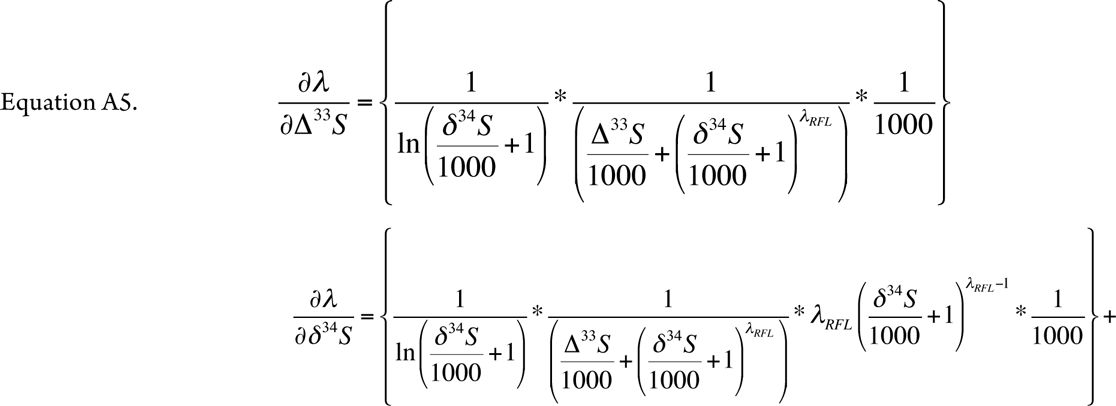

and

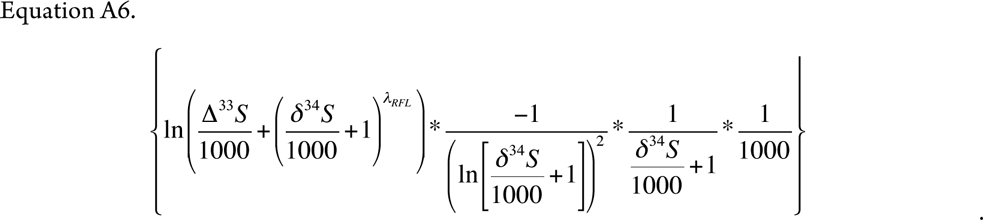

As noted above, the error on lambda σ_l_is dependent on the δ^34^S and δ^33^S. The subscript *RFL* represents the reference fractionation line, which for ^33^λ is 0.515, and for ^36^λ is 1.90. Data for δ^36^S are not discussed in the text, as they yield the same conclusions as δ^33^S, but are included here in Dataframe S1 (https://github.com/bradleylab/DsrAB_enzyme_models).

In total, this leaves our error estimate a function of the following five variables: ^34^ε, σ_R_, *f*, σ_f_, and α. The final error is not an evenly weighted sum of these variables, and in the case presented here, most heavily influenced by the error in concentration data (σ_f_). A sensitivity analysis (Figure A6) on this exercise demonstrates that the errors in *f* far outweigh the analytical uncertainty in a measurement of *R*, and dominate the magnitude of the final σ_Rf_.

### Appendix 5. Data compilations and statistical analysis

#### A5.1 Compilation and statistical analysis of pure-culture MSR fractions

To place our DsrAB enzyme-specific fractionation factor in context with the previous 65 years of pure-culture experimental work, we compile all available observations from studies using axenic cultures of MSR (Figure 4), in the following experimental systems: *batch* (closed-system, in vivo, whole-cell), *chemostat* (open-system, in vivo, whole-cell), *resting* (closed-system, in vivo, whole-cell, not growing), *cell-free* (closed-system, ex vivo crude cell extracts, not growing). From these four types of experiments we further subdivide experiments into where sulfate was reduced to sulfide or sulfite was reduced to sulfide. We count each experimental determination (^34^ε_r-p_) and compile them all in the supplemental data files (http://dx.doi.org/10.6084/m9.figshare.1436115), from experiments where less than 10% of the reactant S-species was consumed. Herein we calculate and present column statistics (box-whisker plots in Figure 4) using *Prism5c* (GraphPad, San Diego, CA). The key finding here is that the majority of the means from each set of experiments is significantly less than the previous estimates for the fractionation factor associated with DsrAB (25 to 53‰ (Harrison and Thode, 1957; Rees, 1973; Brunner and Bernasconi, 2005; Farquhar et al., 2003; Johnston et al., 2007)), and that the mean values from all 650+ experimental determinations, regardless of experiment type or whether it was a sulfate-sulfide or sulfite-sulfide experiments, the grand mean for ^34^ε_*r-p*_ falls at 17.9‰ (median at 16.1‰), with the 25^th^ and 75^th^ percentile’s falling at 10‰ and 22.5‰, respectively (Figure 5) – these are all well within the maximum fractionation accounted for by the sum of our DsrAB value (15.3‰) and our literature derived range for sulfate reduction to sulfite (∼4 to 15‰), for a total of 19.3 to 30.3‰ (*see* main text).

#### A5.2 Literature estimates of fractionation during sulfate activation to sulfite

The upstream kinetic isotope fractionation, the result of enzyme mediated sulfate/sulfite exchange in cell-free extract experiments, is between 4 and 15 ‰ (compilation files: http://dx.doi.org/10.6084/m9.figshare.1436115). The mean of these experiments is ^34^ε_SO4/SO3_ = 9.5‰, CI_95%_ = 7.2 to 11.9‰, with and *n* = 12 (column statistics are also permanently available at: http://dx.doi.org/10.6084/m9.figshare.1436115) (Harrison and Thode, 1958; Kaplan and Rittenberg, 1964; Kemp and Thode, 1968; Ford, 1957). Deconvolving this aggregated fractionation factor (^34^ε_SO4/SO3_) in vitro is a target for future pure enzyme experiments focusing on the constituent steps (enzyme specific ^34^ε), as well as the minor isotope fractionations associate with each (*i.e.* ^33^λ’s).

These values represent the fractionation across the sum of the steps incorporating sulfate activation to APS and its concomitant reduction to sulfite (Figure 1). It is important to note that these values were determined using crude-cell extracts, rather than purified enzymes, and not measured over a range of reaction progress (as in Figure 2). Further, available data do not allow for the evaluation of mass balance closure, as we have done here for DsrAB. Given our present understanding of the enzymes involved in this process (Bradley et al., 2011; Pereira et al., 2011), sulfate transport into the cytoplasm followed by activation to APS (Sat) are not likely to directly impact S-isotope compositions, whereas the reduction of APS (APSr) most likely does, due to the breaking of a S-O bond. The sum of transport, Sat and APSr fractionations sit immediately upstream of the DsrAB. Both of these constraints (^34^ε_SO4/SO3_ and ^34^ε_DsrAB_) are interpreted in the context of the MSR data compiled from the literature, which includes lab experiments, natural waters and sediments, as discussed in the main text (Figure 5).

#### A5.3 Statistical analysis of laboratory chemostat and marine sediment fractionations

To apply the compiled sedimentary sulfate reduction rates from Goldhaber & Kaplan (Goldhaber and Kaplan, 1975), we re-plot their log-scale values to a linear scaling (Figure 5) and apply the same non-linear regression one-phase decay model (Y=(Y_0_ – plateau)e^(–kx)^ + plateau) from our recent work on fractionation—rate relationships in MSR (Leavitt et al., 2013), minimizing variance to arrive at the following parameters: *Y*_0_ = 73‰, plateau = 17.3‰, and a decay-constant (*K*) of 6.4 (Figure 5). For the chemostat (open-system) MSR data in the study where we derived this regression model (Leavitt et al., 2013), we re-scale the cell-specific MSR rates to basic volumetric fluxes by multiplying out the number of cells at each sampling point, using the chemostat values from our recent study (Leavitt et al., 2013). Applying the same one-phase decay model and minimize variance, we calculate the following parameters: *Y*_*0*_ = 56.5‰, plateau = 17.3‰, and a decay-constant (*K*) of 0.054. All regressions were calculated using *Prism5c* (GraphPad, San Diego, CA).

### APPENDIX FIGURES

**Figure A1.**
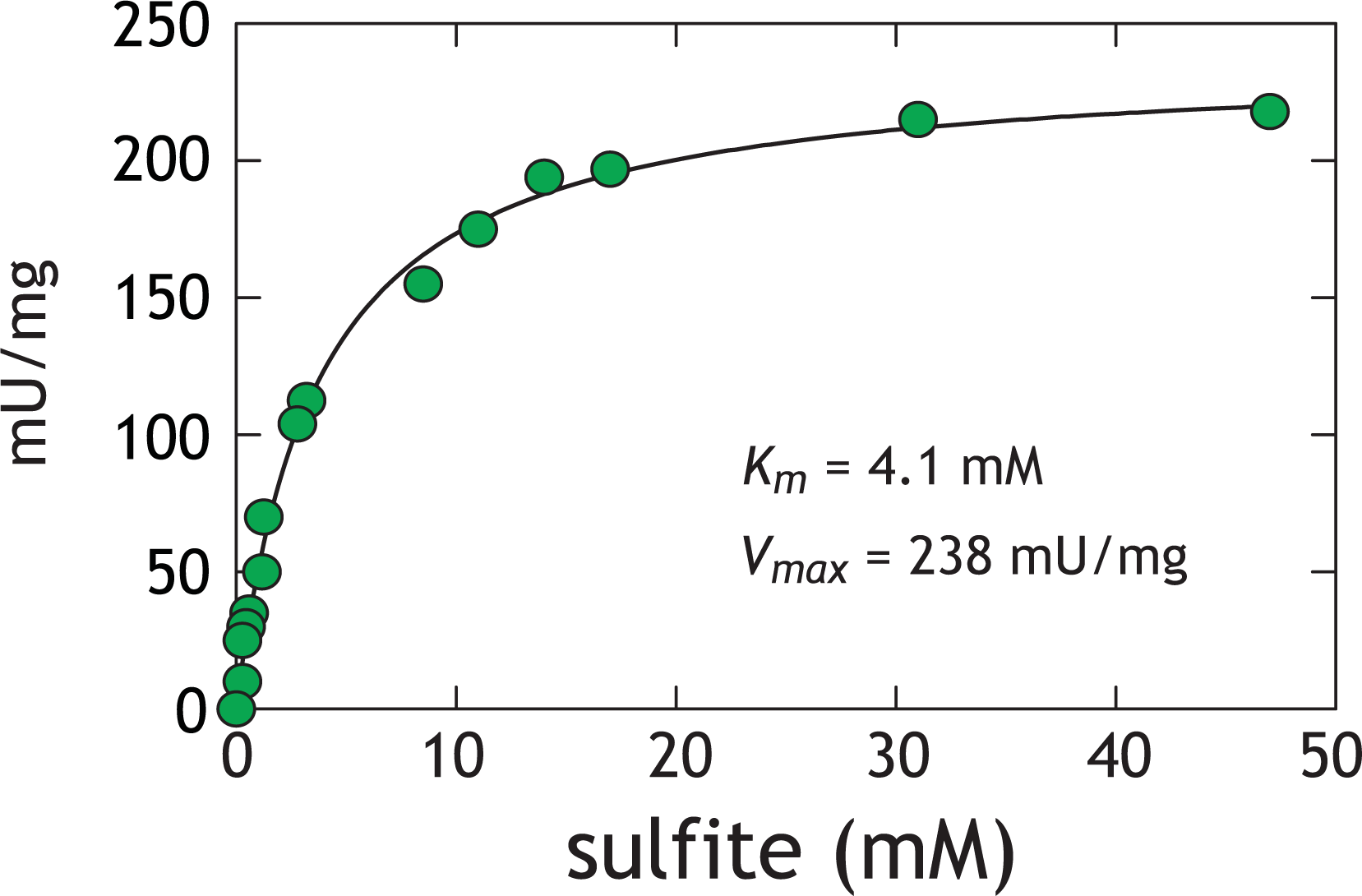
Activity assays with *D. vulgaris* DsrAB. The difference between initial and final concentration of sulfite after two hours was used to calculate the rate. The Michaelis-Menten equation 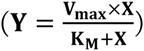 was solved for experimental K_m_ and V_max_under our conditions. The analytical error is less than the size of the symbols (2σ = 1μM). One unit (U) is defined as the quantity of enzyme that catalyzes the conversion of one micromol of substrate per minute. At both 10 and 15mM initial sulfite we are assured to be well above the apparent DsrAB *K*_*m*_ for sulfite. Reaction inhibition was not observed at sulfite concentrations as high as 50 mM (Soriano and Cowan, 1995; Wolfe et al., 1994).

**Figure A2.**
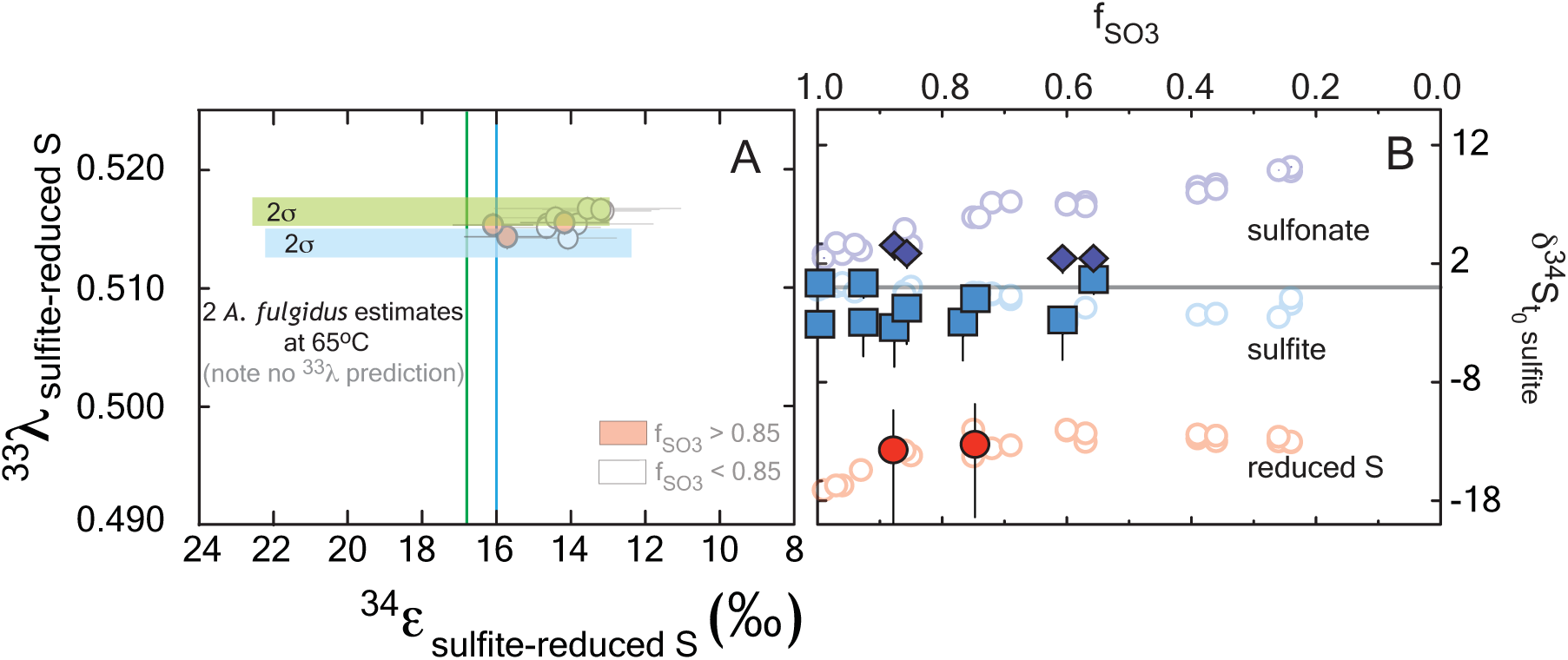
The major isotope ratios (δ^34^S) reported relative to the composition of sulfite at t_0_, for the *A. fulgidus* (closed symbols) and *D. vulgaris* (open symbols) experiments both as a function of reaction progress. The samples sets show a general consistency, particularly at the reduced S sites, despite the significant offet in temperature (20-30**°**C for *D. vugaris* relative to 65**°**C for *A. fulgidus*), consistent with kinetic theory (Bigeleisen and Wolfsberg, 1958), where temperature should impart a minimal effect over this range. The asymmetric error bars on reduced S moieties are a function of the non-linear correction for small sample sizes available for isotope ratio measurements (*see* Appendix text for details).

**Figure A3.**
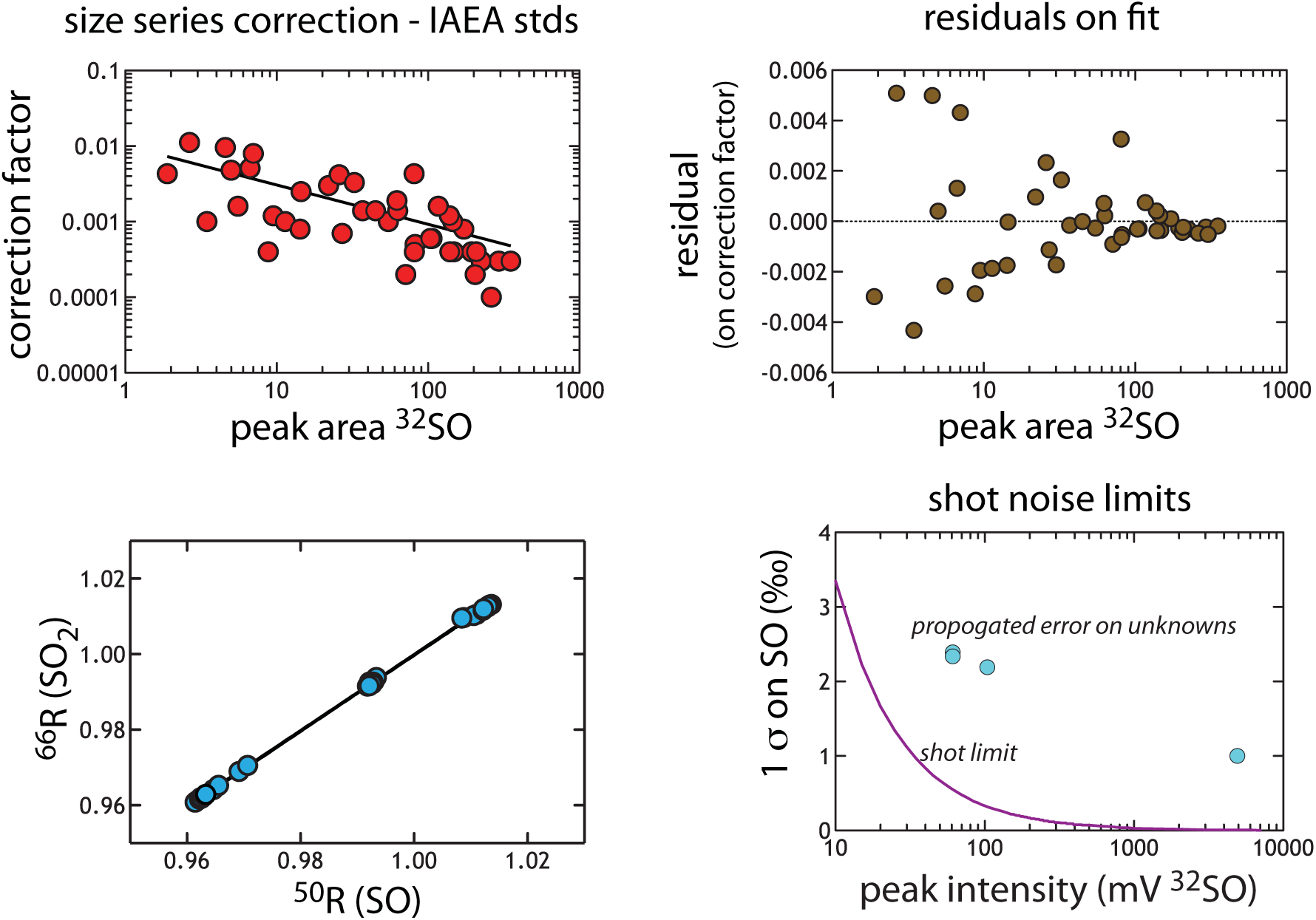
The size series correction calculated from IAEA standards (n = 44). (A) Plots the correction (value from 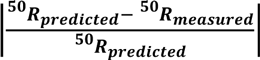 against peak-integrated area). (A) Shows the calculated residual around that fit, demonstrating a symmetric distribution that scales with peak area. This is an illustration of the goodness of the fit. (C) The regression used to convert SO data to an SO_2_ scale. (D) The calculated shot noise for SO as a function of signal intensity (peak height in mV). This precision limit is below that which we propagate through the correction, and is provided for reference here.

**Figure A4.**
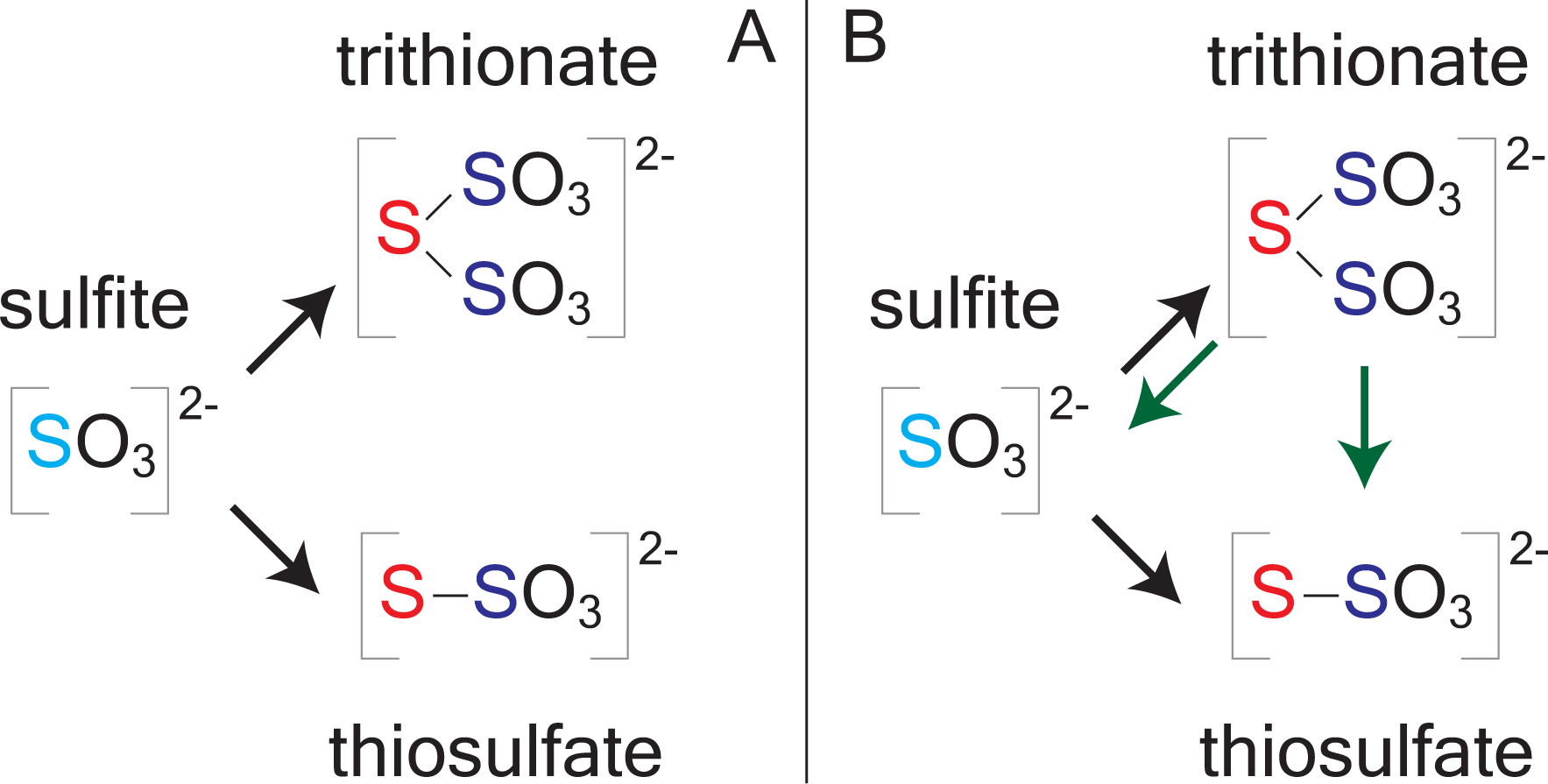
Reaction topology for closed-system models. (A) The simple model for early in the experiments (*f* > 0.85). (B) Shows a more complex reaction model that may be applicable later in the closed-system experiments (*f* < 0.85).

**Figure A5.**
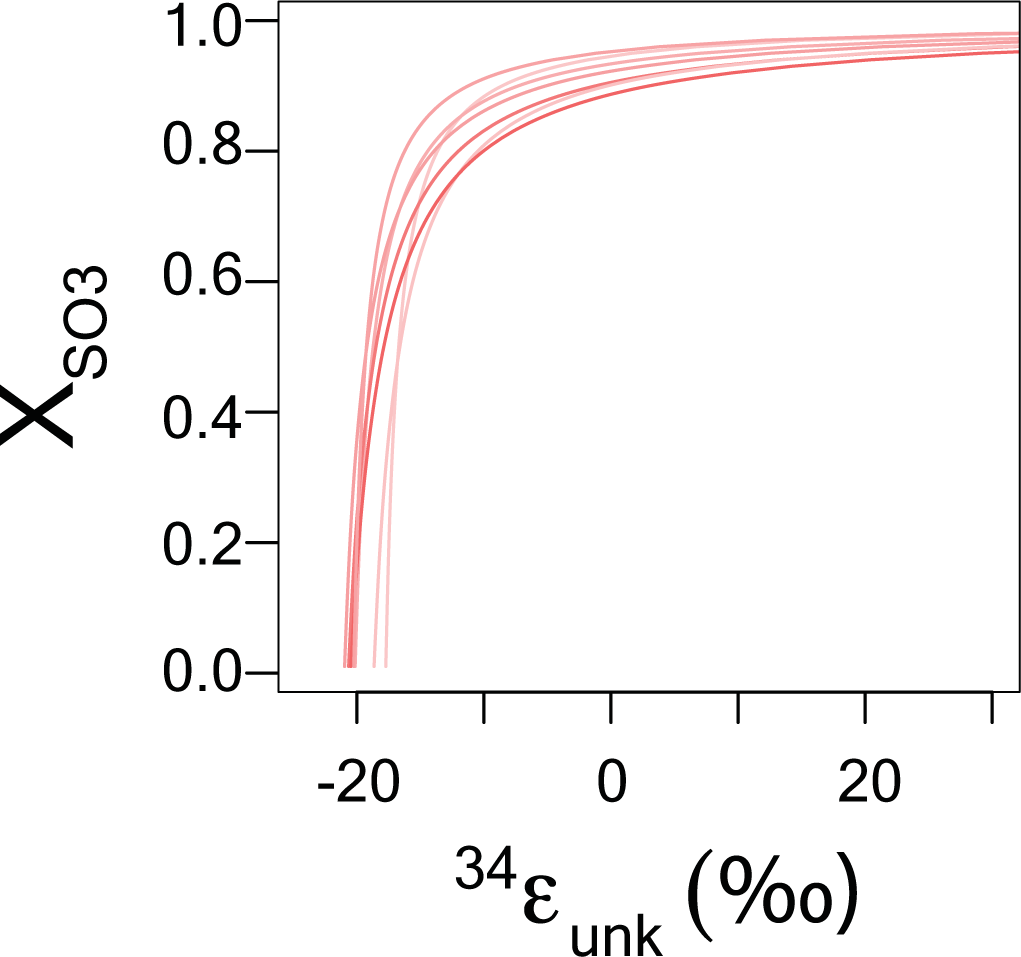
The relationship between the magnitude of a secondary fractionation (^34^α_secondary_) and the proportion of reduced sulfur deriving directly from sulfite reduction (X_SO3_). The balance (1 − X_SO3_) is from the parallel reduction of bound sulfonate. Errors estimates are from the propagation calculation, incorporating both isotope and concentration analysis analytical errors (see below).

**Figure A6.**
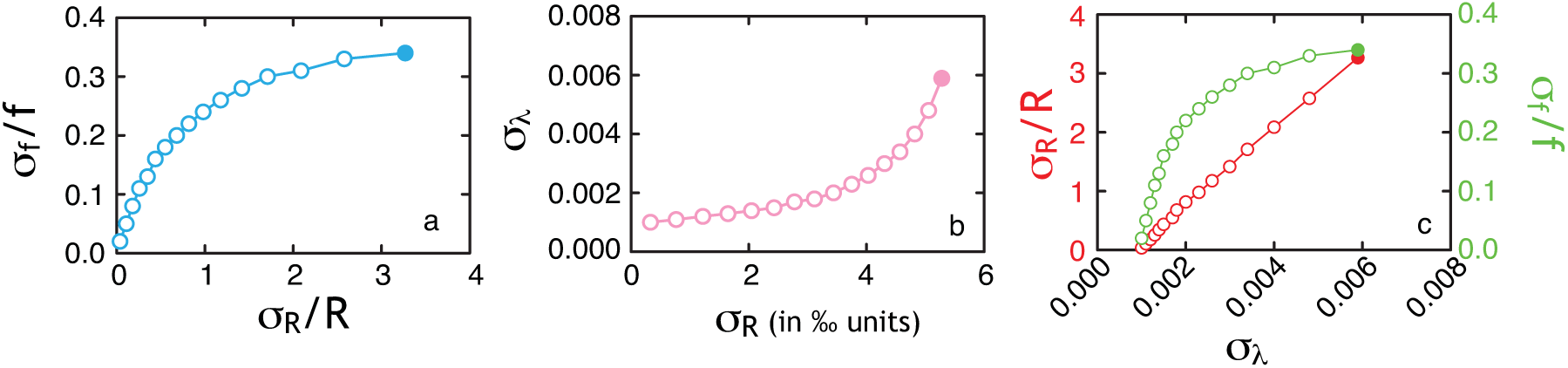
A series of frames (A-C) describing the origin and sensitivity of the controls on the resultant error in δ^34^S and ^33^λ plotted in Figure A2 and 3. For this analysis, ^34^ε is set to 15.3‰ to be consistent with the reductive branch of the DsrAB experiments, the analytical precision of an isotope measurement is 0.2‰ in δ^34^S and 0.008‰ in δ^33^S, and *f* is allowed to vary from 0.6 to 0.9 in 0.02 increments (noted by circles). This range in *f* is chosen to reflect the experimental range, and the circle is filled where *f* = 0.9. In (a), the covariance of the relative error in *f* (σ_*f*_/*f*) is show to correlate with the error in the relative isotope ratio, σ_*R*_/*R*. As the calculated error in the *R* derived from the Rayleigh equation (σ_*R*_) increases, the consequence is an increase in the error in ^33^λ (b). Finally, the relationship between the errors in both *f* and *R*, as they contribute to the error on ^33^λ, are presented in frame c. Frames b and c do not approach the origin as a result of *f* not approaching the limits of 0 and 1, and also due to the multivariate nature of the propagation.

